# Evidence for flexible regulation of movement and decision vigor in a reward-oriented task

**DOI:** 10.1101/2025.11.25.690375

**Authors:** Adrien Conessa, Arnaud Boutin, Bastien Berret

## Abstract

The vigor of movement and decision-making is fundamental to many reward-oriented behaviors. While some theories propose that movement and decision are jointly invigorated to maximize a global utility (e.g., mixing reward, effort, and time), alternative perspectives suggest that the brain can separately invigorate them when advantageous. We tested these competing hypotheses using a foraging-like task that allowed experimentally assessing the vigor of movement (reaching to reward location) and that of decision-making (harvesting reward). By using a block-wise design, we measured the effects of independent manipulations of time and effort on reach and harvest durations. Decoupled effects were found on reach and harvest durations, with no inter-individual consistency between them. A model allowing separate, yet inter-dependent, optimizations of movement and decision vigor based on distinct time costs sensitive to recent temporal history predicted these results more accurately than a global utility model. These findings indicate that movement and decision vigor may flexibly depend on whether shared versus distinct time costs underlie behavior; co-regulation would arise when the vigor of movement and decision is governed by a shared time cost but decoupling may emerge when relevant for the task.

**Significance statement:** Coordinating movement and decision-making is fundamental to many reward-oriented behaviors. Current theories suggest that the brain may invigorate behavior by co-regulating movement and decision, but some experimental findings rather point to a decoupling. To shed new light on this debate, we developed a foraging-like task promoting independent and comparable manipulation of time and effort. We show that movement and decision are mostly decoupled in this task, invigorated by distinct time-cost signals. Previously experienced delays tend to amplify these time costs, thereby exerting a strong influence on behavior and indirectly linking movement and decision. Co-regulation or decoupling may hinge on whether movement and decision vigor relies on shared or distinct time-cost signals.

## Introduction

Most of our daily behaviors are influenced by the time and effort they require, as well as the rewards they are expected to yield. For instance, picture yourself at a crowded conference buffet. What dictates how fast you approach the buffet and how long you forage before relocating? Depending on your time constraints, the difficulty of obtaining food amidst the crowd, and your current level of hunger, your behavior may vary significantly. There is evidence that time, effort, and reward are represented in various brain regions, in particular in the ventral striatum and basal ganglia circuitry as a whole (Frank et al., 2007; Prévost et al., 2010; Robbe and Dudman, 2020; Salamone et al., 2003). Neuroeconomic models have been developed to account for the complex interplay among these factors and their influence on volitional behaviors. A prominent model posits that the times allocated to movement and decision originate from a common control mechanism maximizing a behavioral utility, namely the global capture rate, defined as the value of all rewards minus all efforts divided by total time (Shadmehr et al., 2016; Yoon et al., 2018).

The *common control hypothesis* (Shadmehr et al., 2016) relies on a single, unifying formula to simultaneously predict both movement and decision times, which necessarily implies that these two critical behavioral variables are strongly and rigidly intertwined. For example, an alteration in either movement effort or movement time should entail a change in decision time (and vice versa), ensuring the maximization of the global capture rate. Several works involving eye saccades, arm reaching or the Tokens task have provided support for this theoretical model (Sukumar et al., 2024; Thura et al., 2014; Yoon et al., 2018). However, the incomplete experimental validation of the model’s predictions prompted some studies to argue instead for a *separate control hypothesis* for movement and decision vigor (see Thura et al., 2025 for a review). For example, it was found that an increase in movement time leads to faster decisions (Saleri Lunazzi et al., 2021), or that an increase in decision and movement effort respectively leads to a decrease in movement time (Yoon et al., 2018), without affecting the amount of reward collected (Sukumar et al., 2024). However, separate control does not imply that the processes are strictly independent, as recent behavioral history may influence the current invigoration. These deviations from the predictions of the common-utility model may partly reflect differences in experimental paradigms across studies. They may also stem from variations in time and effort manipulations, as well as from the inherent difficulty of dissociating their influence on goal-directed behaviors (Morel et al., 2017; Saleri Lunazzi et al., 2021; Verdel et al., 2023)—despite evidence for their distinct evaluation in brain subsystems (Prévost et al., 2010).

To test the *common* versus *separate* control hypotheses of behavioral vigor, we therefore designed a foraging-like task that introduced comparable manipulations of time and effort in both movement and decision processes. Similar to the example introduced earlier, the task involved two sequential phases: participants first had to reach a location (the movement phase), and then harvest a reward by remaining there until they decided to relocate (the decision phase). Importantly, to balance time and effort in movement and decision phases, we leveraged a robotic interface and considered an isometric task in a well-controlled environment. The task was designed such that time and effort in each phase were of comparable magnitudes, and isolated by independent manipulation of time through the addition of temporal delays. We then tested the predictions of the common-utility model, including the consistency of inter-individual differences during movement and decision: individuals who reach faster should also spent less time harvesting (Reppert, 2025). In contrast, if movement and decision are separately controlled, their durations should exhibit relative independence and a lack of inter-individual consistency. This outcome would support an alternative separate-control model where the movement and decision phases are optimized individually, allowing for flexible decoupling of their invigoration.

## Methods

### Participants

A total of forty-four subjects participated to this experiment. Twenty healthy volunteers (8 females, mean age: 26.3 ± 4.7 years) were recruited by local advertisements for the main experiment. Additionally, fourteen healthy volunteers (8 females, mean age: 27.2 ± 3.0 years) and ten healthy volunteers (5 females, mean age: 27.1 ± 3.3 years) were respectively recruited for the first and second control experiments. All participants met the following inclusion criteria: aged between 18 and 35 years, without any psychiatric and/or neurological disorder, without any impaired function of the upper limbs or uncorrected visual impairment. The Local Ethics Committee from the Université Paris-Saclay (CER-Paris-Saclay-2024-24) approved the experimental protocol, which conformed to relevant guidelines and regulations. All participants gave written informed consent before inclusion.

### Experimental design

#### Main experiment

Participants sat on a chair at a distance of 50 cm in front of a computer screen (Fig. 1B). Participants were connected with their right hand (irrespective of hand dominance) to an HRX-1 robotic wrist exoskeleton (Human Robotix) controlled at 100 Hz, which included a torque sensor. The robotic handle was fixed by a hardware lock, allowing only isometric contraction while measuring the torque exerted by the participant. The maximal torque produced by each participant during a maximal voluntary contraction (*τ_MVC_*, in Nm) was evaluated to calibrate the required effort during the task. Electromyography (EMG) of the right flexor carpi ulnaris and extensor carpi radialis muscles were also recorded. To assess the effect of time and effort in a reward-oriented behavior, we developed a foraging-like task in which participants pushed against a robotic exoskeleton using the back of their hand. A red dot (representing the participant’s position) and an empty green patch (representing the target location) were respectively displayed on the left and right sides of the screen, separated by a distance of 40 cm (Fig. 1C, D). The task consisted of two distinct phases: a reach phase and a harvest phase.

**Figure 1.**
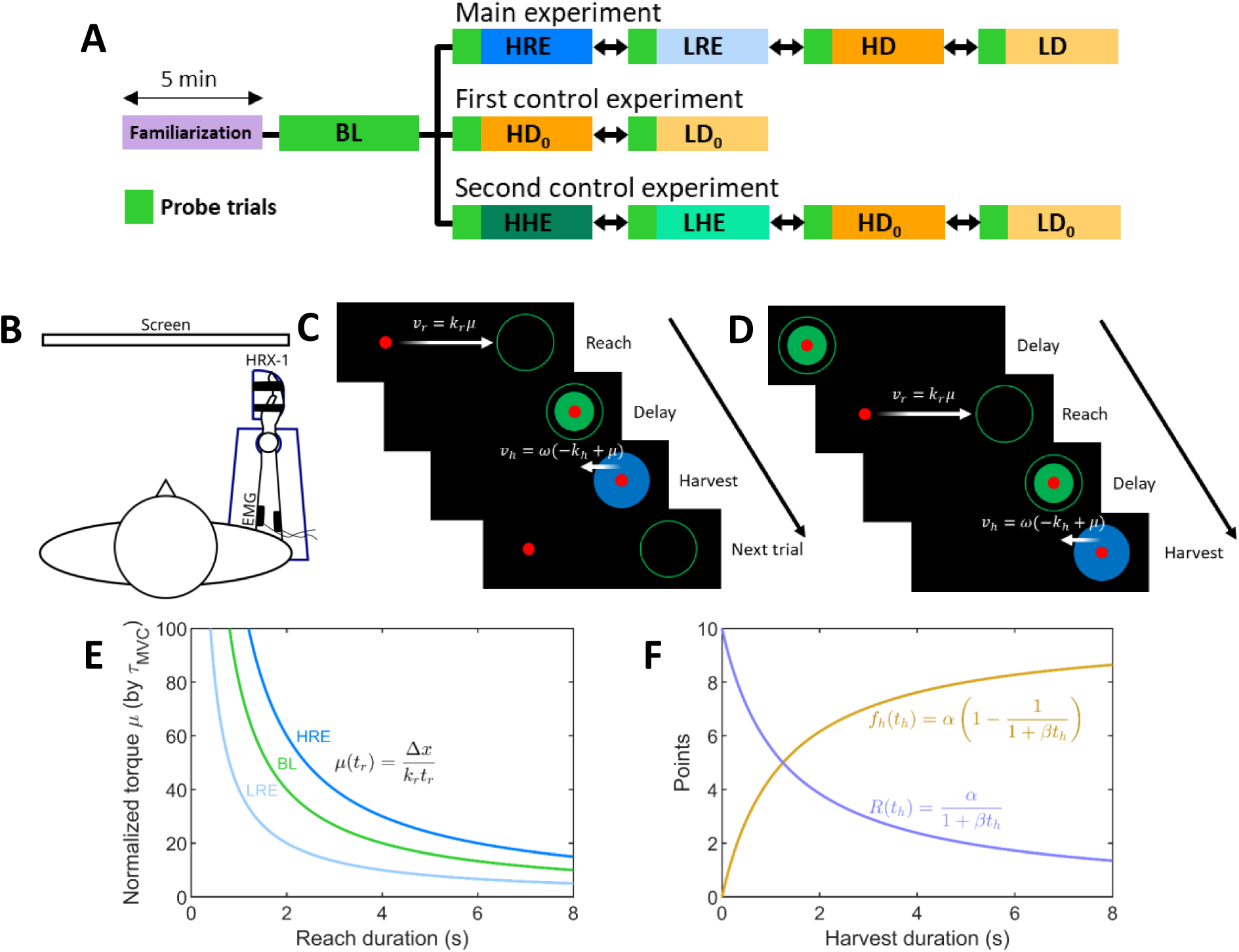
Experimental design. **(A)** Organization of the different experiments. Each experiment began with a familiarization block, followed by a baseline block. In the main and second control experiments, participants completed four additional conditions, whereas in the first control experiment, only two additional conditions were tested. The order of conditions (excluding the familiarization and baseline blocks) was randomized across participants. To minimize potential after-effects, the first 5 trials of each block were “wash-out” trials, using the same parameters as the baseline condition. (**B**) Illustration of the experimental setup. Participants were connected to an HRX-1 robotic exoskeleton (in blue) in front of a computer screen. Two EMG electrodes were placed over the wrist flexor and extensor muscle groups. **(C)** Example of one trial from the main experiment. The trial began with a reach phase, in which participants moved a red dot with a velocity proportional to the applied torque. Once reaching the patch, a delay elapsed during which the patch gradually filled. This was followed by the harvest phase, during which participants resisted a virtual force field to stay within the patch and collect points. Pressing the space bar ended the trial, resetting the red dot to the starting position and initiating the next trial. **(D)** Example of one trial from the control experiments. A delay period was introduced before the reach phase, during which a target gradually emptied over time. **(E)** Relationship between reach duration and the average normalized torque applied for each reach effort condition. **(F)** The harvest function (in yellow) depicts the number of points collected over time within a patch. This function is subject to maximization. In terms of minimization, the equivalent function is the ‘cost of reward’ (in blue), obtained by negating *f* and adding the maximum reward per patch, *α* (see modeling section below). BL: Baseline; HRE: High Reach Effort; LRE: Low Reach Effort; HD: High Delay; LD: Low Delay; HHE: High Harvest Effort; LHE: Low Harvest Effort.

During the reach phase, participants were required to push against the robotic exoskeleton to move the red dot toward the green patch. A direct relationship between the normalized torque *μ* (unitless, as normalized by *τ_MVC_*) applied by the participant against the robot and the red dot velocity *v_r_* (in m/s) was defined through Eq. 1, as follows:

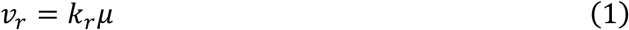

The coefficient *k_r_* (in m/s) defined the maximal motion speed achieved when producing the maximal torque. For instance, for 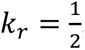, a torque of 20%*τ_MVC_* was required to move with a speed of 10 cm/s, which would in turn correspond to a reach duration of 4 seconds if the motion were at constant speed (Fig. 1E). Here, participants were instructed to move at a self-selected speed, while avoiding jerks or pauses during the movement. Once they reached the empty green patch, the latter began to fill gradually within 3 seconds, delaying the onset of the subsequent harvest phase. Participants were instructed simply to stay inside the patch during this delay (denoted by *td* later on), and they had no isometric torque to produce. Once fully filled, the patch turned blue, marking the beginning of the harvest phase. In this phase, participants were able to collect points by staying within the patch, withstanding a virtual force field that pushed the red dot toward the left edge of the patch. To simulate this force field, a constant leftward velocity was added to the dot velocity, such that the cursor’s velocity during harvest was set as follows:

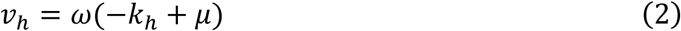

where *v*_*h*_ is the dot velocity (in m/s), *ω* is a conversion coefficient 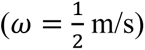, *k*_*h*_ (unitless) is the strength of the virtual force field, and *μ* is the normalized torque defined above. To counteract the leftward velocity and stay inside the patch (i.e., *v*_*h*_ = 0 m/s), participants had to generate a torque proportional to *τ_MVC_* by the factor *k*_*h*_ (e.g., if *k*_*h*_ = 0.20, a torque of 20%*τ_MVC_* was required to withstand the virtual force field). The normalized torque *μ* required to withstand the virtual force field is thus simply given by solving Eq. 2, that is *μ* = *k*_*h*_.

The number of points collected within a patch increased with harvest duration *t*_*h*_ and was specified by Eq. 3 to account for the patch depletion (see also Fig. 1F) similarly to other studies (Sukumar et al., 2024; Yoon et al., 2018):

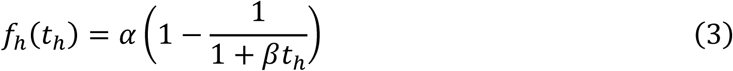

In this function, *α* represents the maximum reward that can be obtained from each patch and *β* represents the rate of patch depletion. As a result, the longer participants remained within the same patch, the fewer points they earned per second. The number of points collected from each patch was added to a cumulative score counter. Participants were free to stay within a patch for as long as they wanted to collect more reward, or to move on to the next patch (i.e., the next trial). To do so, they simply had to press the space bar with their left hand, which deactivated the virtual force field and caused the red dot to relocate on the left side of the screen, alongside a new empty green patch on the right side.

Participants were instructed to maximize the number of points collected during each 5-min block, while remaining free to adopt the pace they considered best suited to them. The parameter *α* was fixed at 10 points, and the value of *β* was selected to ensure that reach and harvest durations could be of similar magnitudes. The selection procedure was as follows: assuming a steady strategy throughout a block with reach duration *tr* and patch-filling delay *td* over the 5-min block, the optimal harvest duration 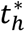 to maximize the total number of points harvested can be computed and is given by 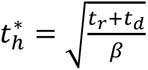. By setting *β* = 0.8, the optimal harvest duration would be close to 3 seconds for a reach duration and a delay of 3 seconds each. Importantly, this setting allows us to design an experiment with potentially comparable durations between the reach and harvest phases, avoiding very unbalanced phase durations that could affect their relative importance in the task.

Participants were first required to perform a 5-min familiarization block of the task. During this block, a cumulative score counter was displayed at the top center of the screen. They also benefited from feedback about the number of points harvested from the current patch, displayed just above it. To ensure that participants understood the instruction to maximize the number of points and produced harvest durations that comply with it, feedback on the optimal harvest duration for each patch was displayed based on the average reach durations of the 3 last trials. The text displaying the current patch’s harvested points gradually changed color according to the ongoing harvest duration, shifting from blue (too short), to gold (optimal), and finally to red (too long). Following the familiarization phase, participants performed a series of 5 test blocks separated by 1-min rest periods (Fig. 1A). During these blocks, all feedback was removed, leaving only the red dot and the target patches visible. As a result, participants received no information about their cumulative score, the number of points collected during the current harvest, or the optimal harvest duration. Different conditions in terms of effort and delay were tested to analyze if and how participants changed their behavior.

The first of the five blocks served as a baseline (BL) condition, in which the effort and delay parameters corresponded to those used during the familiarization phase (3-sec delay, 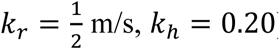). The main experiment comprised four additional blocks, in each of which a single parameter was independently manipulated. In the High Reach Effort (HRE) condition, reach effort was increased such that a torque of 30%*τ_MVC_* resulted in a red dot velocity of 10 cm/s (i.e., with 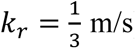) (Fig. 1E). Conversely, in the Low Reach Effort (LRE) condition, the required torque was reduced to 10%*τ_MVC_* for the same velocity (i.e., with *k_r_* = 1 m/s) (Fig. 1F). In the High Delay (HD) condition, the delay between the reach and harvest phases was extended to 5 seconds, while in the Low Delay (LD) condition, this delay was shortened to 1 second. The order of these 4 blocks was randomized across participants. Also, to prevent any after-effect between conditions, the first five trials of each block replicated the baseline condition settings (20%*τ_MVC_* and 3-sec delays). Participants were aware of these “wash-out” trials at the beginning of each block and were told that something would change just after, although the specific nature of the change was not explicitly told. Finally, at the end of each block they were asked to estimate the number of points collected, and to guess what aspect of the task had changed compared to the wash-out trials. This procedure allowed us to assess participants’ explicit awareness of the manipulations in effort or time constraints.

#### Control experiments

To gain a deeper understanding of how time and effort influence reward-oriented behavior, two additional control experiments were conducted. First, it can be hypothesized that the delay introduced in the main experiment may have a sequential effect, primarily influencing the subsequent phase (specifically, the harvest phase). To investigate how the timing of delays affects reach and harvest durations, both control experiments used a similar design to the main experiment, but included an additional delay before the reach phase. At the beginning of each trial in the control experiments, the red dot appeared within a green filled target located on the left side of the screen, which gradually emptied over a default 3-second delay. Once completely emptied, the target disappeared, and a new empty green patch appeared on the right side of the screen, 40 cm away, as in the main experiment. Thus, each trial was composed of four sequential components: an initial delay, the reach phase, a second delay (kept constant at 3 seconds for all conditions of the control experiments), and the harvest phase.

In the first control experiment, participants completed a familiarization block, a baseline block, and two additional experimental conditions. In the High Delay (HD_0_) condition, the initial delay before reaching was extended to 5 seconds, while in the Low Delay (LD_0_) condition, it was reduced to 1 second.

In the second control experiment, participants again completed a familiarization block, a baseline block, the HD_0_ and LD_0_ conditions, and two additional experimental conditions with changes in term of harvest effort. In the High Harvest Effort (HHE) condition, the required effort to counter the virtual force field of the harvest phase was increased such that participants needed to apply a torque of 30%*τ_MVC_* to stay within the patch. In the Low Harvest Effort (LHE) condition, this requirement was reduced to 10%*τ_MVC_*.

In sum, by combining the main experiment and the control ones, we were able to thoroughly study how variations in terms of effort and time across the different phases affect such reward-oriented behaviors.

#### Behavioral metrics

For all experiments, the behavioral analyses aimed at assessing how time and effort modulations influenced reach and harvest durations, the two dependent variables in this task. In the main experiment, reach duration was defined as the time elapsed from the onset of the trial to the moment the red dot entered the empty green patch (i.e., the start of the inter-phase delay). In the control experiments, reach duration was defined as the interval between the end of the initial delay and the entry of the red dot into the empty green patch (i.e., the start of the inter-phase delay). It is noteworthy that reach duration also encompasses reaction time. The cyclic nature of this paradigm does not foster the specific study of reaction times. For example, the reach phase is preceded by the harvest phase in the main experiment, which leads to no clear stopping point (without clear drop in velocity) between the two. Similarly, participants could easily anticipate target appearance in the control experiments, and initiate their movement before the actual reach onset. Nevertheless, for the sake of completeness, additional analyses were conducted to estimate ‘reaction times’ using a conservative approach, and reported in Supplementary Information (Fig. S2). Finally, for all experiments, harvest duration was defined as the time elapsed between the end of the inter-phase delay and the instant participants pressed the space bar to switch to the next trial. The median reach and harvest durations, and the interquartile range (IQR), were computed for each condition, without including the “wash-out” trials.

Stimuli presentation and response registration were controlled using the MATLAB R2021b software from The MathWorks (Natick, MA) and the Psychophysics Toolbox extensions.

### EMG data acquisition and pre-processing

Electromyographic activity was recorded using two electrodes (Cometa MiniWave) placed over the flexor carpi ulnaris and extensor carpi radialis muscles, following SENIAM guidelines for electrode placement (Hermens et al., 2000). Prior to electrode application, the skin was shaved and cleaned, and electrodes were positioned 2 cm apart along the muscle belly by applying a conductive patch. Data were recorded using at a 1-kHz sampling rate. The raw EMG signal was filtered using a bandpass filter with cutoff frequencies of 20 Hz and 450 Hz (two-pass finite impulse response [FIR] bandpass filter). The EMG signal was then normalized to the maximal absolute EMG value obtained during three maximal voluntary contraction. To estimate muscle effort, the integral of the absolute normalized EMG signal was computed (see Fig. S4 for additional information).

### Modeling of time-effort effects on reach and harvest durations

We investigated how time and effort are integrated by the central nervous system (CNS) to determine: (i) the duration allocated to moving from an initial position to a rewarding patch and (ii) the duration allocated to collecting reward within the patch. The common-utility model posits that the behavioral goal of participants is to maximize the global capture rate *J̅* defined as the total reward harvested across all patches minus the total effort expended divided by the total time (Shadmehr et al., 2016; Yoon et al., 2018). A major feature of the global capture rate is that the effort and time spent in the reach and harvest phases are taken into account simultaneously to determine optimal reach and harvest durations (Fig. 2A). This model also assumes knowledge about the environment, which is coherent with the block-wise design of our experiment. Hence, we aimed to test the theoretical predictions of the common-utility model in the present task.

**Figure 2.**
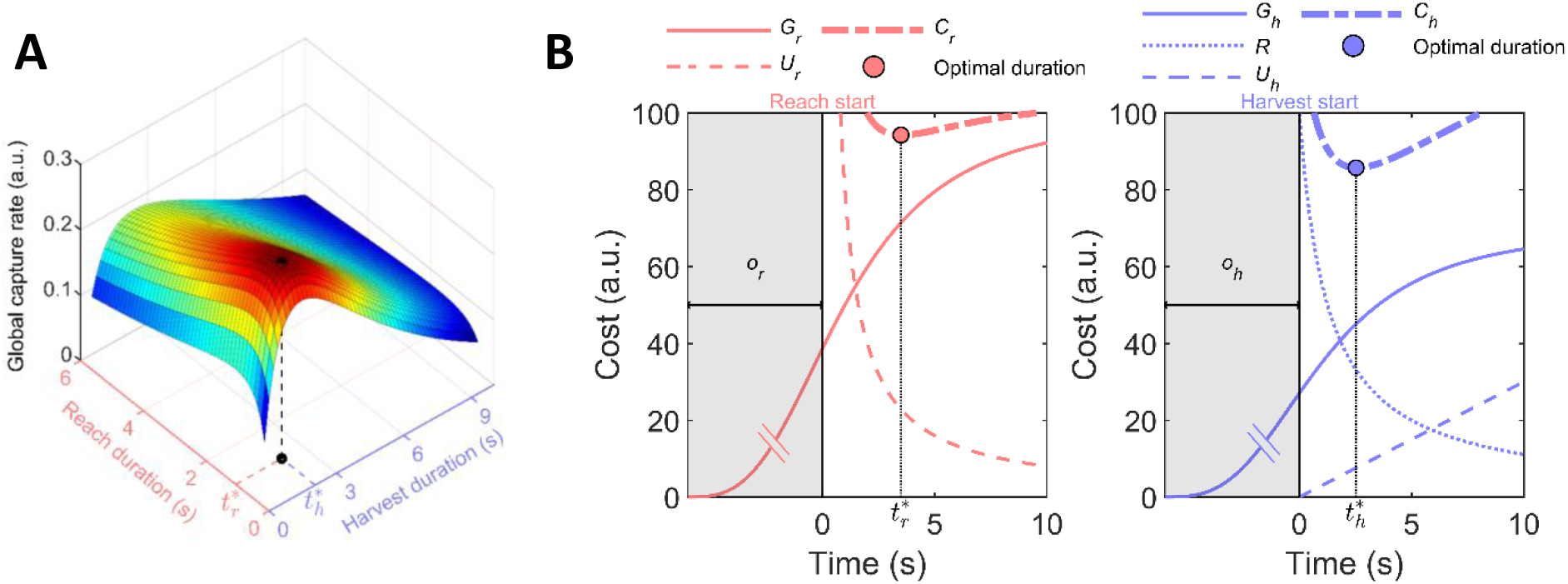
Common-utility and separate-cost models. **(A)** Utility as a function of the reach and harvest durations. In this utility-based framework, reach and harvest phases are jointly regulated, with optimal durations selected to maximize a single time-discounted utility function that integrates effort, reward, and time. **(B)** An alternative model is proposed such that reach (red, left panel) and harvest (blue, right panel) are optimized separately. Here, optimal durations are selected to minimize distinct subjective cost functions (*C_r_* and *C*_*h*_; thick dot-dashed lines), each comprising at least a cost of time (*G_r_* and *G*_*h*_; dashed lines) and a cost that decreases over time (*U_r_* and *R*; solid lines). Note that the cost of time integrates all preceding delays, implying mutual influence of one phase over another depending on their onset order. Here, the general case is considered without any assumption on the phase order, but see supplementary information (Fig. S6 and Supplementary Video) for representation of a typical trial of the experiments illustrating the inter-dependence induced by the offset. *U*_*h*_: harvest’s cost of effort; o_r_: reach time offset.

#### Common-utility model

The utility *J* of a single trial was defined as the reward harvested, minus reach/harvest efforts, divided by total time. We already represented the reward harvested by the function *f*_*h*_(*t*_*h*_) as it is defined by the task (Eq. 3). In addition, we assume that reach effort *U_r_*(*t_r_*) and harvest effort *U*_*h*_(*t*_*h*_) are respectively functions of reach duration *t_r_* and harvest duration *t*_*h*_, which specially makes sense in our isometric settings. Finally, a temporal discounting can be incorporated to discount the utility with the passage of time, via the term *G*(*t*_*h*_, *t_r_*). Altogether, the temporally-discounted utility of a trial (or local capture rate) can be expressed as:

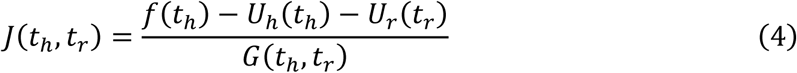

According to the optimal foraging theory, the goal of participants would be to maximize the global capture rate over N patches, rather than optimizing the local capture rate for each patch independently (Yoon et al., 2018). This approach is well-suited for dynamic environments, where patch rewards, efforts, and delays vary across trials. However, in our experimental design, the environment within each block was fixed, meaning that maximizing the discounted utility of a single, repeated trial effectively maximizes the global capture rate. Consequently, by working on average durations, we can compute an average temporally-discounted utility per trial across the 5-min block that should be maximized. To reduce the influence of outlier trials, we computed and analyzed the median of each behavioral metric. We aimed at predicting the optimal median harvest and reach durations, respectively 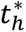 and 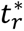, from the maximization of *J*. The derivatives of *J* with respect to *t*_*h*_ and *t_r_* are given by:

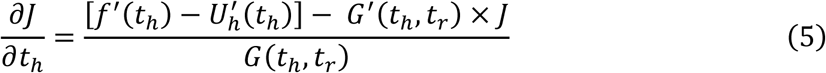

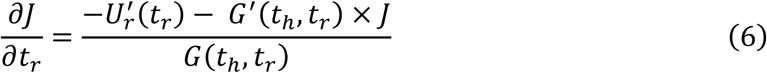

The optimum harvest and reach durations can be found when both of these equalities are simultaneously equal to 0. The next step is to define the functions *U_r_*(*t_r_*), *U*_*h*_(*t*_*h*_) and *G*(*t*_*h*_, *t_r_*) in our task.

Cost of effort during reaching (*U_r_*): in our experimental design, the velocity is linearly proportional to the normalized torque applied, *v* = *k_r_μ* where *k_r_* is a scaling coefficient (in m/s). The distance Δ*x* covered during a reach of duration *t_r_* is given by 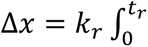. Here, Δ*x* is fixed at 40 cm throughout our experiment, and *k_r_* remains constant within each block, implying that 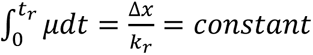. Let 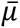 be the average normalized torque applied by the participant during the reach phase, the integral part (the torque-time integral) is then equal to 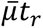, leading to 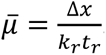. This can be considered as a first estimation of the effort cost of the reach phase (Shadmehr et al., 2016). Yet, it was shown that during isometric contractions, the transient period before producing a steady force can be costlier than maintaining that force, at least for durations of less than two seconds (Russ et al., 2002). Accordingly, in addition to the torque-time integral, the work and the torque-rate may contribute to the overall metabolic cost of isometric contraction (van der Zee and Kuo, 2021). The work is equal to 0 here (as no actual movement occurs), and the energetic cost induced per unit of time by the torque-rate can be considered proportional to 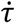 (or 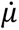 if normalized by *τ_MVC_*) (van der Zee and Kuo, 2021). Hence, the total torque-rate cost *E* can be characterized by the integral of 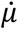 between *t*_0_ and *t_max_*, the time at which maximum normalized torque *μ_max_* is attained and maintained. Consequently, if we start at 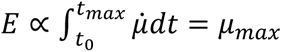. The torque-rate cost was restricted from *t*_0_ to *t_max_* because the remaining duration is mainly characterized by null or negative 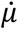, which reflects muscle relaxation in an isometric paradigm, and may marginally affect the energetic cost. As linear mixed model analyses (*lme4* package in R) showed that average normalized torque (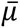) is a good predictor of maximum normalized torque (*μ_max_*) for each condition (see Fig. S7), 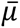 thus reflects both the torque-time integral and the torque-rate costs in our experimental design. Therefore, we consider a simple cost of effort proportional to the average torque applied during the reach phase. Hence, we defined *U_r_* = *b*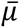 where *b* is a free weighting parameter, allowing us to express *U_r_* as a function of *t_r_*:

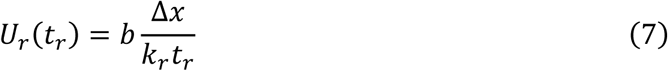

where Δ*x* is fixed and *k_r_* depends on the experimental condition. The cost of reaching is illustrated in Fig. 1E for different *k_r_*.

Cost of effort during harvest (*U*_*h*_): the function *U*_*h*_(*t*_*h*_) represents the effort exerted by participants to resist the constant virtual force field during the harvest phase. As for the reach phase, the torque-rate and the torque-time integral should be taken into account to estimate the energy cost. Again, the torque-time integral is equal to 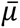*t*_*h*_, and 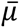 should be inherently close to *k*_*h*_ to counteract the virtual force field (see Eq. 2). Hence, the torque-time integral is proportional to *k*_*h*_*t* _*h*_. For the torque rate, we previously showed that *E* ∝ *μ_max_*, with *μ_max_* conveniently representing the torque required to withstand the virtual force field, and thus *μ_max_* ≈ *k*_*h*_ ≈ 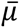. Altogether, we defined a simple cost of effort for the harvest phase, which increases linearly with time and is proportional to both the torque-rate and the torque-time integral via the weighting parameter *a*:

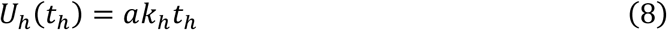

Temporal discounting function (*G*): the function *G*(*t*_*h*_, *t_r_*) was defined to account for all durations (*t*_*h*_, *t_r_*) and delays (*t_d_*), weighted by a parameter γ that determines the rate of discounting (Shadmehr et al., 2016):

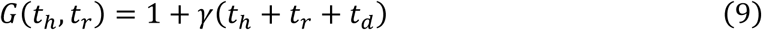

Higher values of the γ parameter produce steeper temporal discounting of the behavioral utility, allowing the model to capture individual differences in discount rates (Jimura et al., 2009). Thus, the temporally-discounted utility of one trial in our experiment can be described by a function depending on the three free parameters *a*, *b* and *γ*:

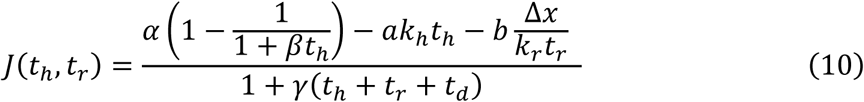

where the other parameters are fixed within each block (i.e., *α*, *β*, Δ*x*, *k_r_*, *k*_*h*_).

Using this definition of the utility function and Eqs. 5 and 6, we aimed to find the best set of parameters that could predict the participants’ median reach and harvest durations across conditions. The Sequential Least Squares Programming (SLSQP) optimization algorithm with constraints (SciPy library) was chosen to find this solution. A loss function was defined to minimize the residual sum of squared errors (RSS) between the predicted and observed median reach and harvest durations for each condition. The constraints were defined such that, for each condition, 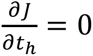 and 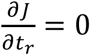 (Eqs. 5 and 6). It is noteworthy that the three free parameters were shared across all conditions within an experiment. The boundaries for parameters *a* and *b* were set between 0 and 100, and between 0 and 5 for the parameter *γ*. Given that the convergence of the local optimization algorithm depends on the initial parameter estimates, 100 optimizations with a random initial guess were performed, and only the best-fitting parameters were retained.

#### Separate-cost model

An alternative model assuming a separate control of movement and decision vigor was considered (Fievez et al., 2024; Reynaud et al., 2020). This alternative model also aims to predict the participants’ reach and harvest durations but through separate optimization problems (Fig. 2B). In this model, reach and harvest durations are selected in order to balance costs specifically relevant to reaching and harvesting, each comprising systematically two components: a cost decreasing with phase duration and a cost increasing with phase duration. A key component of this model is to assume that the brain plans behaviors by considering a cost incurred by the passage of time, termed cost of time (Berret and Jean, 2016; Shadmehr, 2010). This cost can be balanced with the other costs in our task (e.g., cost of effort and cost of reward for reaching and harvesting, respectively). In principle, such a cost of time may also integrate or be sensitive to delays preceding each phase (Constantino and Daw, 2015; Haith et al., 2012). It is noteworthy that this specific property implies that both optimizations are separated, albeit not strictly independent. Reach and harvest may interact with each other through changes in their duration, which would delay the onset of the consecutive phase and be integrated as a temporal offset in the cost of time underlying the current vigor choice. In the motor domain, a sigmoidal shape has been found for the time cost in a range of reach durations from 500 to 1500 milliseconds (Berret et al., 2018; Berret and Jean, 2016). As sigmoid function can locally capture on this scale a wide range of shape (concave upward/downward, linear), we consider the relatively large family of sigmoidal functions for the cost of time as follows:

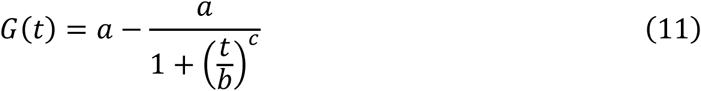

where *a*, *b* and *c* are free parameters: *a* is the upper asymptote, *b* is the half-max point location and *c* is the relative steepness of the curve. If *c* is greater than 1, there is an inflexion point where the cost of time switches between a fast exponential-like increase to a slow hyperbolic-like increase. The time *t_i_* at which the inflection point happens is given by 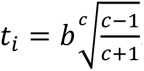. The time *t* in Eq. 11 is the total time elapsed from the *beginning* of the trial and is not restricted to the reach and harvest durations, but accounts for all preceding durations and delays of the current trial. Given the highly cyclical nature of the required behavior in our 5-minute blocks, we make the assumption that this cost of time resets at the beginning of each trial. Hence, for the reach phase, *G* would be evaluated at *t* = *t_r_* + *o_r_*, where *o_r_* is an offset for the reach phase, and for the harvest phase, at *t* = *t*_*h*_ + *o*_*h*_, where *o*_*h*_ is an offset for the harvest phase. These offsets integrate all delays and durations preceding the corresponding phase within the trial. Unlike previous cost of time models, this model thus considers that the cost of time could be offset by the delays recently experienced by participants. Note that the time cost may be specific to each phase: its main property is to characterize the participants’ sensitivity to elapsed time in each phase. It may reflect a growing urgency signal (Thura et al., 2025), which, depending on the context, be alternatively interpreted as a boredom signal (Berret et al., 2018) or implicit motivation in the task (Mazzoni et al., 2007).

Regarding the costs decreasing with phase duration, they are objectively distinct for the two phases. For the reach phase, the latter can be represented as the physical cost of effort *U_r_* (Eq. 7). Regarding the harvest phase, the physical cost of effort *U*_*h*_ (Eq. 8) does not fulfil this requirement, as it increases with time. A third cost must be included, termed here the cost of reward, *R*(*t*_*h*_), that prevents the participant from leaving the patch early. This cost directly arises from the harvest function (Eq. 3), such that *R*(*t*_*h*_) = *α* − *f*_*h*_(*t*_*h*_) (Fig. 1F). Hence, the cost of reward decreases as the harvest progresses, with the cost being maximal when *t*_*h*_ = 0, giving us the following equation:

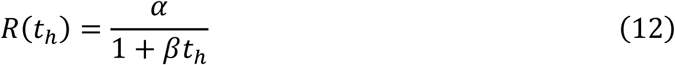

To summarize, we can define a cost for reaching (*C_r_*; Eq. 13) and a cost for harvesting (*C*_*h*_; Eq. 14) in our experimental task, as follows:

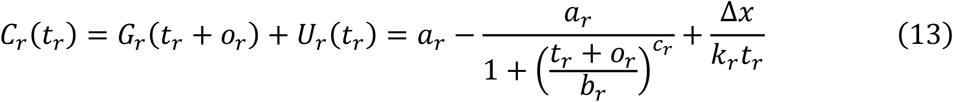

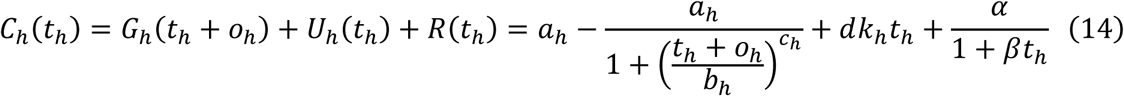

where *a_r_*, *b_r_* and *c_r_* are the free parameters of the reach-related cost of time, *a_h_*, *b_h_* and *c_h_* are the free parameters of the harvest-related cost of time, and *d* is a free weighting parameter. Let *t_d_* be the duration of all delays preceding the corresponding phase; then *o_r_* is an offset for the reach phase such that here *o_r_* = *t_d_*, and *o*_*h*_ is an offset for the harvest phase such that here *o*_*h*_ = *t_d_* + *t_r_*, where *t_r_* is the preceding reach duration (set to the experimental value for the model fitting). In this model, reach and harvest durations would be chosen separately by the CNS to minimize these respective costs, given previously experienced durations and based on a similar mechanism balancing time-increasing and time-decreasing costs (Fig. 2B). Interactions between reach and harvest durations are thus only indirect, via the cost of time offsets. Optimal durations are then found by solving 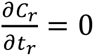 and 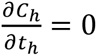.

As for the common-utility model, we used the SLSQP optimization algorithm with constraints (SciPy library) to determine the best set of parameters that could predict the participants’ median reach and harvest durations across conditions. A driver function was defined to minimize the RSS between the predicted and observed median durations and the optimization constraints were set such that 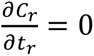 and 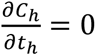 for each condition. This model required six free parameters to find the optimal reach and harvest durations (three for each phase). Importantly, the same parameter values were shared across all conditions within the experiment. The lower limit for all parameters was set at 0, the upper limits for *a*, *b*, and *c* were respectively 1000, 100, 10 for both the reach and harvest costs of time, and the upper limit for *d* was set at 100. As with the common-utility model, the optimization procedure was run 100 times with a different initial random guess, and only the best-fitting parameters were retained.

### Statistical analysis

All subjects were included in statistical analyses. Nonparametric statistical analyses were performed to comply with the sample sizes of all experiments. Pairwise comparisons between conditions were assessed using Wilcoxon signed-rank tests, and relationships between variables were evaluated using Spearman correlations. All tests were two-tailed, with a significance threshold set at *p* < 0.05. Effect size for the Wilcoxon signed-rank test was reported using the Rank-Biserial correlation (r_rb_). Where appropriate, multiple comparisons were corrected using the Benjamini-Hochberg procedure (Benjamini and Hochberg, 1995). To compare the performance of the common and separate models, we computed the Akaike Information Criterion (AIC) using the following formula: 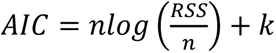 where n is the number of samples, RSS is the residual sum of squared errors, and k is the number of free parameters plus one (Burnham and Anderson, 2004). Consequently, lower AIC values indicate better model performance.

## Results

### Main experiment

Wilcoxon signed-rank tests were performed to assess the effect of changes in terms of reach effort (HRE vs LRE) on reach and harvest durations (Fig. 3A, B). Regarding reach duration, the analysis revealed a significant effect (W = 210, Z = 3.92, *p* < 0.001, r_rb_ = 1), highlighting an increase in durations for the HRE condition (*t_r_* = 3.35 s, IQR = 1.31 s) compared to the LRE condition (*t_r_* = 1.65 s, IQR = 0.61 s). Regarding harvest duration, the analysis failed to detect a significant effect (W = 131, Z = 0.97, *p* = 0.35, r_rb_ = 0.25), suggesting quite marginal differences between the HRE (*t*_*h*_ = 3.54 s, IQR = 1.55 s) and the LRE (*t*_*h*_ = 3.25 s, IQR = 1.43 s) conditions.

**Figure 3.**
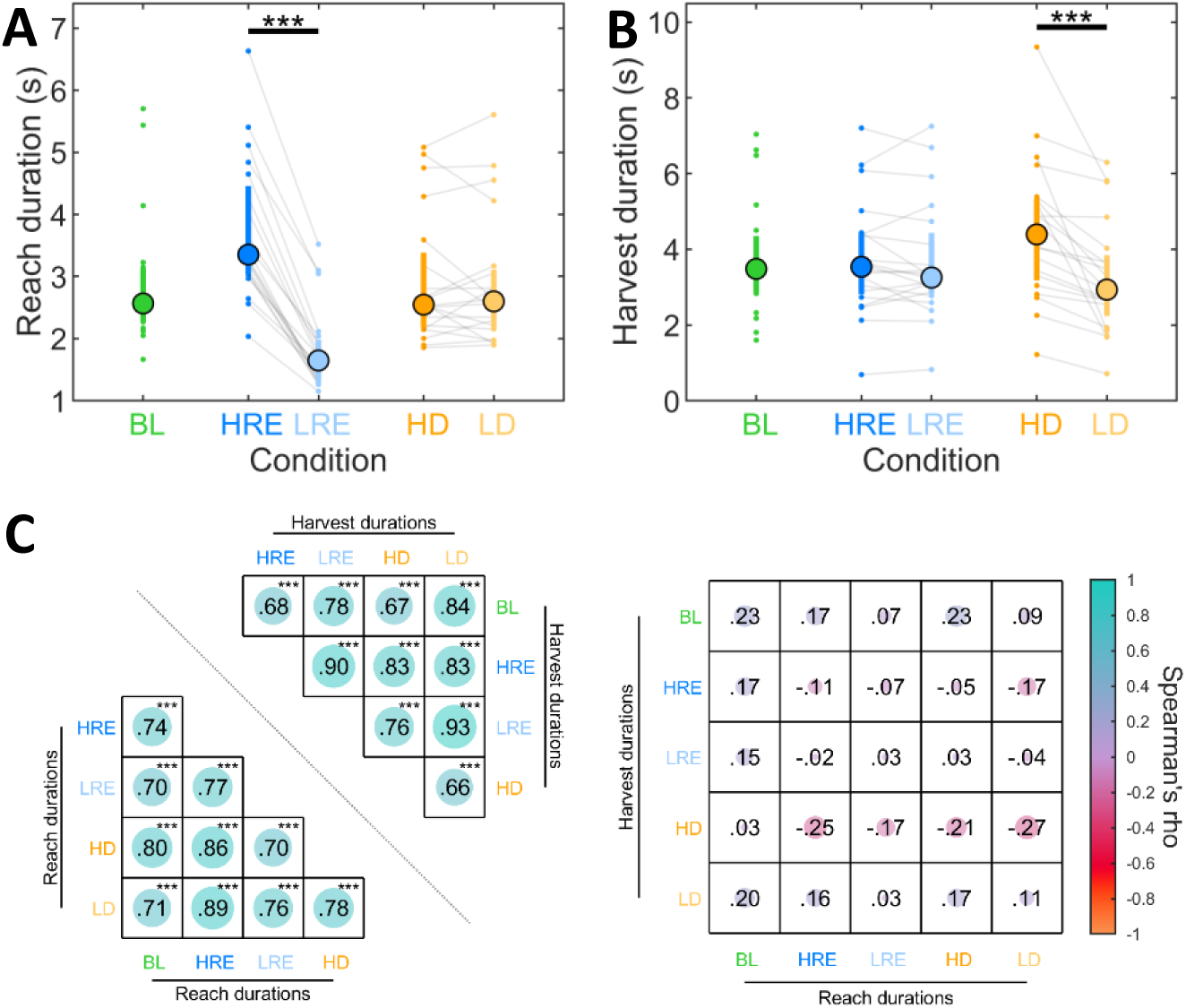
Effects of time and reach effort on reach and harvest durations (main experiment). Reach **(A)** and harvest durations **(B)** for each condition of the main experiment. Colored circles represent median durations, and bars represent interquartile ranges (n = 20 for each condition). Individual data points are displayed as small colored dots. Thin grey lines connect compared observations across conditions. The stars represent p-values associated with Wilcoxon signed-rank tests. **(C)** Left panel: heatmap of Spearman correlation coefficients between all pairs of condition for reach (lower triangle) and harvest (upper triangle) durations. Right panel: heatmap of Spearman correlation coefficients between reach and harvest durations for each pair of conditions. The stars represent p-values associated with the analyses, after correction for multiple comparisons using the Benjamini-Hochberg procedure (Ntests = 45). *** *p* < 0.001. BL: Baseline (green); HRE: High Reach Effort (blue); LRE: Low Reach Effort (light blue); HD: High Delay (orange); LD: Low Delay (yellow).

Wilcoxon signed-rank tests were performed to assess the effect of changes in terms of delays (HD vs LD) on reach and harvest durations (Fig. 3A, B). Regarding reach duration, the analysis failed to detect a significant effect (W = 99.0, Z = -0.22, *p* =0.84, r_rb_ = -0.06), suggesting quite marginal differences between the HD (*t_r_* = 2.55 s, IQR = 1.08 s) and the LD (*t_r_* = 2.60 s, IQR = 0.88 s) conditions. Regarding harvest duration, the analysis revealed a significant effect (W = 210, Z = 3.92, *p* < 0.001, r_rb_ = 1), highlighting an increase in durations for the HD condition (*t*_*h*_ = 4.39 s, IQR = 2.08 s) compared to the LD condition (*t*_*h*_ = 2.94 s, IQR = 1.36 s). Trial-by-trial presentation of reach and harvest durations are provided in Supplementary information, confirming the stability of these behavioral effects (Fig. S1 A).

Then, relationships between reach and harvest durations were investigated at the inter-individual level to assess the consistency of participants’ behaviors. Spearman correlations were carried out on reach durations between pairs of conditions and reported in a heatmap (Fig 3C, lower triangle). The analyses show a robust inter-individual consistency during reaching: the fastest participants in one condition were also the fastest in another. Similarly, Spearman correlations were carried out on harvest durations between pairs of conditions and reported in a heatmap (Fig 3C, upper triangle). The analyses show robust inter-individual consistency during harvesting: participants who harvested for longer periods in one condition also harvested for longer periods in another. Finally, to assess the relationship between reach and harvest durations, additional Spearman correlations were performed across all condition pairs and presented in a separate heatmap (Fig. 3C, right panel). Interestingly, no significant relationship was found, indicating that participants’ reaching speed was not predictive of their harvest duration; reach and harvest durations tended to be unrelated at the inter-individual level. Note that all correlations were corrected for multiple comparisons using the Benjamini-Hochberg procedure to control for the false discovery rate (Ntests = 45). Finally, we also sought to verify whether participants who showed the largest difference (in percentage) in terms of reach or harvest duration between one pair of conditions (HRE-LRE and HD-LD) also showed the largest difference between the other pair of conditions. Pairwise correlation analyses revealed no significant relationship, indicating that participants who exhibited large reach-duration differences (HRE-LRE) did not necessarily show large harvest-duration differences (HD-LD).

### First control experiment

A first control experiment was conducted with 14 additional participants to determine whether the effects of delay observed in the main experiment depend on the onset timing within the trial. The experimental design was similar to the baseline condition of the main experiment, except that an additional delay was introduced prior to the reach phase.

Wilcoxon signed-rank tests were performed to assess the effect of changes in terms of delays appearing before the reach phase (HD_0_ vs LD_0_) on reach and harvest durations (Fig. 4A, B). Regarding reach duration, the analysis revealed a significant effect (W = 85, Z = 2.04, *p* =0.042, r_rb_ = 0.62), highlighting an increase in reach durations for the HD_0_ condition (*t_r_* = 2.11 s, IQR = 0.85 s) compared to the LD_0_ condition (*t_r_* = 1.99 s, IQR = 0.69 s). Regarding harvest duration, the analysis revealed a significant effect (W = 97.0, Z = 2.79, *p* = 0.003, r_rb_ = 0.85), highlighting an increase in harvest durations for the HD_0_ condition (*t*_*h*_ = 4.87 s, IQR = 1.88 s) compared to the LD_0_ condition (*t*_*h*_ = 4.37 s, IQR = 2.32 s). Trial-by-trial presentation of reach and harvest durations are also available in Supplementary information (Fig. S1 B).

**Figure 4.**
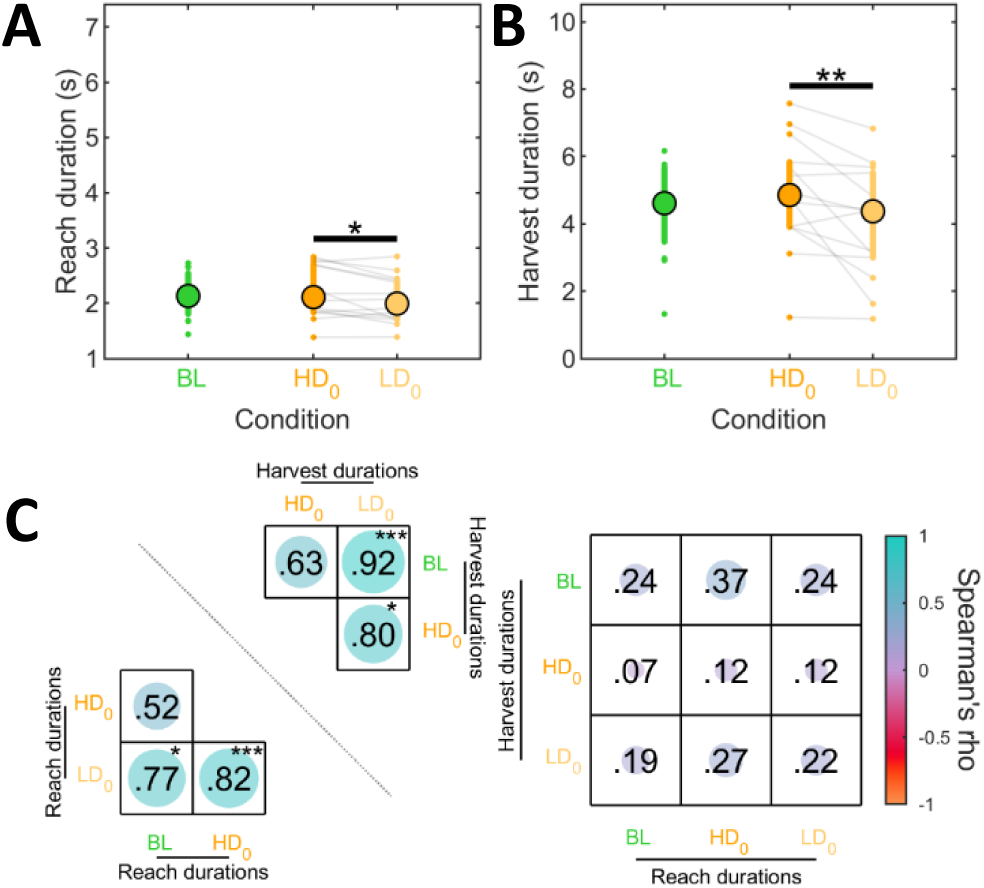
Effects of delay’s timing onset on reach and harvest durations (first control experiment). Reach **(A)** and harvest durations **(B)** for each condition of the first control experiment. Colored circles represent median durations, and bars represent interquartile ranges (n = 14 for each condition). Individual data points are displayed as small colored dots. Thin grey lines connect compared observations across conditions. The stars represent p-values associated with Wilcoxon signed-rank tests. **(C)** Left panel: heatmap of the correlations separately for reach (lower part) and harvest (upper part) durations between each pair of conditions. Right panel: heatmap of the correlations between reach and harvest durations for each pair of conditions. Spearman correlation coefficients (rho) are reported for each correlation. The stars represent p-values associated after correction for multiple comparisons using the Benjamini-Hochberg procedure (Ntests = 15). * *p* < 0.05 ** *p* < 0.01; *** *p* < 0.001. BL: Baseline (green); HD_0_: High Delay (orange); LD_0_: Low Delay (yellow).

Relationships between reach and harvest durations were then investigated at the inter-individual level. Spearman correlations were carried out on reach durations between pairs of conditions and reported in a heatmap (Fig. 4C, lower triangle). The analyses showed a robust inter-individual consistency for reaching: the fastest participants in one condition were also the fastest in another condition, except between BL and HD_0_ conditions (rho = 0.52, *p* = 0.18), where only a positive non-significant trend was observed. Similarly, Spearman correlations were carried out on harvest durations between pairs of conditions and reported in a heatmap (Fig. 4C, upper triangle). The analyses showed robust inter-individual consistency for harvesting: participants who harvested for longer periods in one condition also harvested for longer periods in another condition, except between BL and HD_0_ conditions (rho = 0.63, *p* = 0.05). Finally, to assess the relationship between reach and harvest durations, additional Spearman correlations were performed across all condition pairs and presented in a separate heatmap (Fig. 4C, right panel). Interestingly, no significant relationship was found, indicating that participants’ reaching speed was not predictive of their harvest duration. All correlations were corrected for multiple comparisons using the Benjamini-Hochberg procedure to control for the false discovery rate (Ntests = 15). Finally, similarly to the main experiment, participants who showed the largest differences in terms of reach duration between the HD_0_ and LD_0_ conditions, did not necessarily show the largest differences in harvest duration.

### Second control experiment

A second control experiment was performed on 10 additional participants to assess the effects of the harvest effort and to confirm the effects of before-reach delays on reach and harvest durations. The same experiment as the first control one was performed, but variations in terms of harvest efforts were also introduced.

Wilcoxon signed-rank tests were performed to assess the effects of changes in terms of harvest effort (HHE vs. LHE) on reach and harvest durations (Fig. 5A, B). Regarding reach duration, the analysis failed to detect a significant effect (W = 20, Z = -0.78, *p* = 0.49, r_rb_ = - 0.27), suggesting similar durations between the HHE (*t_r_* = 1.93 s, IQR = 0.55 s) and LHE (*t_r_* = 1.99 s, IQR = 0.74 s) conditions. Regarding harvest duration, the analysis failed to detect a significant effect (W = 32, Z = 0.46, *p* = 0.70, r_rb_ = 0.16), suggesting quite marginal differences between the HHE (*t*_*h*_ = 2.96 s, IQR = 1.23 s) and the LHE (*t*_*h*_ = 2.93 s, IQR = 1.23 s) conditions. Thus, our variations of the harvesting effort did not lead to significantly different reach and harvest durations, despite clear effects in the effort exerted (Fig. S4-5).

**Figure 5.**
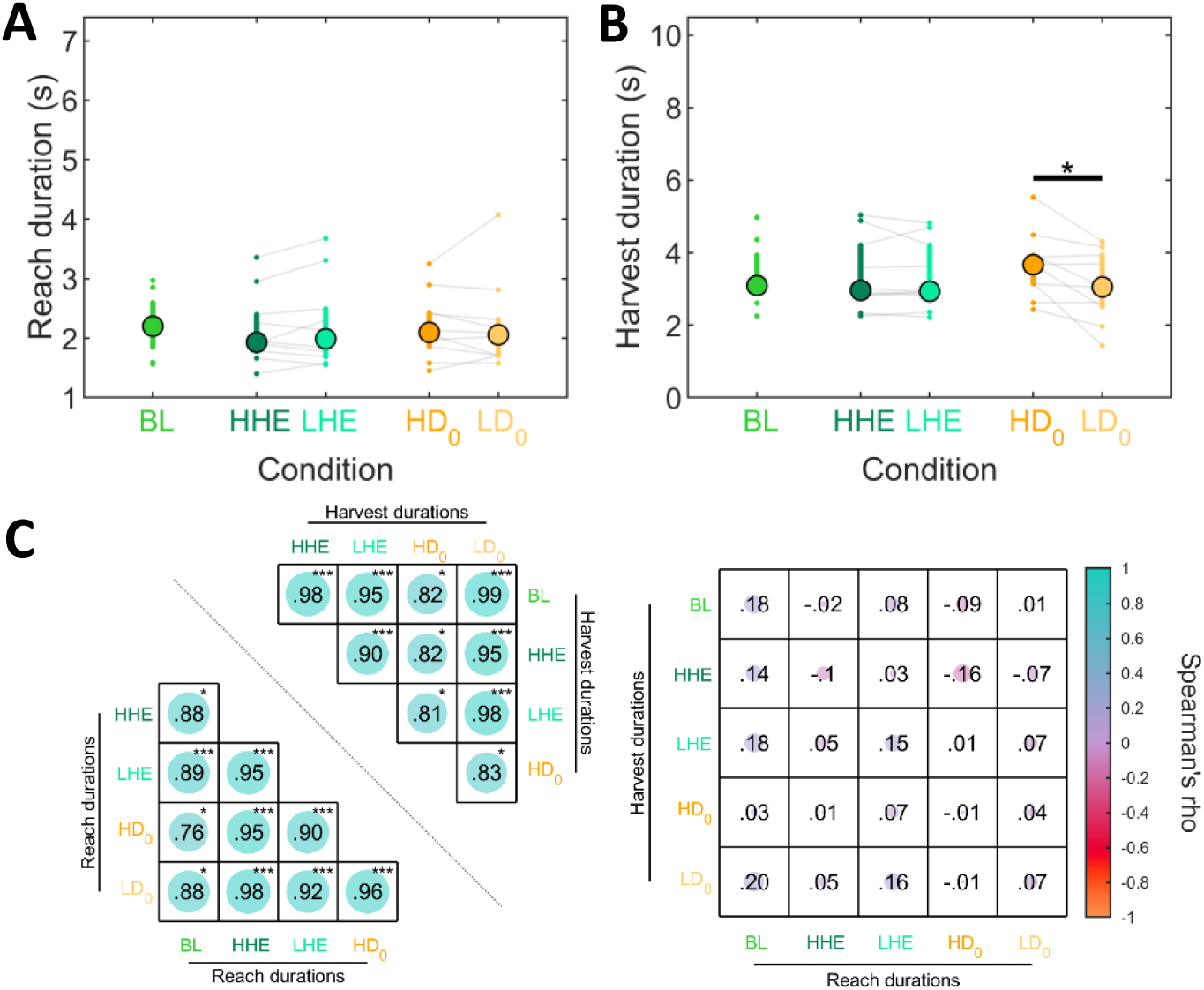
Effects of time and harvest effort on reach and harvest durations (second control experiment). Reach **(A)** and harvest durations **(B)** for each condition of the first control experiment. Colored circles represent median durations, and bars represent interquartile ranges (n = 10 for each condition). Individual data points are displayed as small colored dots. Thin grey lines connect compared observations across conditions. The stars represent p-values associated with Wilcoxon Signed-rank tests. **(C)** Left panel: heatmap of the correlations separately for reach (lower part) and harvest (upper part) durations between each pair of conditions. Right panel: heatmap of the correlations between reach and harvest durations for each pair of conditions. Spearman correlation coefficients (rho) are reported for each correlation. The stars represent p-values associated after correction for multiple comparisons using the Benjamini-Hochberg procedure (Ntests = 45). *** *p* < 0.001. BL: Baseline (green); HHE: High Harvest Effort (dark green); LHE: Low Harvest Effort (light green); HD_0_: High Delay (orange); LD_0_: Low Delay (yellow).

Wilcoxon signed-rank tests were performed to re-assess and eventually confirm the effects of changes in terms of delays (HD_0_ vs LD_0_) on reach and harvest durations (Fig. 5A, B). Regarding reach duration, the analysis failed to detect a significant effect (W = 35, Z = 0.76, *p* = 0.49, r_rb_ = 0.27), suggesting quite marginal differences between the HD_0_ (*t_r_* = 2.10 s, IQR = 0.53 s) and LD_0_ (*t_r_* = 2.06 s, IQR = 0.57 s) conditions. This confirms that the effect observed in the first control experiment on reach duration was indeed relatively marginal. Regarding harvest duration, the analysis confirmed a significant effect (W = 49, Z = 2.19, *p* = 0.027, r_rb_ = 0.78), highlighting an increase in harvest duration for the HD_0_ condition (*t*_*h*_ = 3.67 s, IQR = 0.77 s) compared to the LD_0_ condition (*t*_*h*_ = 3.05 s, IQR = 1.34 s). Trial-by-trial presentation of reach and harvest durations are also available in Supplementary information (Fig. S1 C).

Relationships between reach and harvest durations were then investigated at the inter-individual level. Spearman correlations were carried out on reach durations between pairs of conditions and reported in a heatmap (Fig. 5C, lower part). The analyses showed a robust inter-individual consistency: the fastest participants in one condition were also the fastest in another condition. Similarly, Spearman correlations were carried out on harvest durations between pairs of conditions and reported in a heatmap (Fig. 5C, upper part). The analyses showed robust inter-individual consistency: participants who harvested for longer periods in one condition also harvested for longer periods in another condition. Finally, to assess the relationship between reach and harvest durations, additional Spearman correlations were performed across all condition pairs and presented in a separate heatmap (Fig. 5C, right panel). Interestingly, no significant relationship was found, indicating that participants’ reaching speed was again not predictive of their harvest duration. All correlations were corrected for multiple comparisons using the Benjamini-Hochberg procedure to control for the false discovery rate (Ntests = 45). Finally, similarly to the main experiment, participants who showed the largest differences in terms of reach or harvest durations between one pair of conditions, did not necessarily show the largest differences between the other pair of conditions.

### Model predictions

Having examined how changes in effort and time affect reach and harvest durations in this task, we next assessed how well the common-utility model and the separate-cost model could account for participants’ behavior. The common-utility and separate-cost models have respectively three and six free parameters used to fit the observed data. Hence, only the main experiment and the second control experiment (each providing 10 median duration data points) were used for fitting the models (Fig. 6). Moreover, for fair comparisons, model performance was assessed using the Akaike Information Criterion (AIC) to take into account the greater flexibility of the separate-cost model and provide comparable measure between models (see methods). A summary of models’ fit and parameter estimates is reported in Table 1.

**Figure 6.**
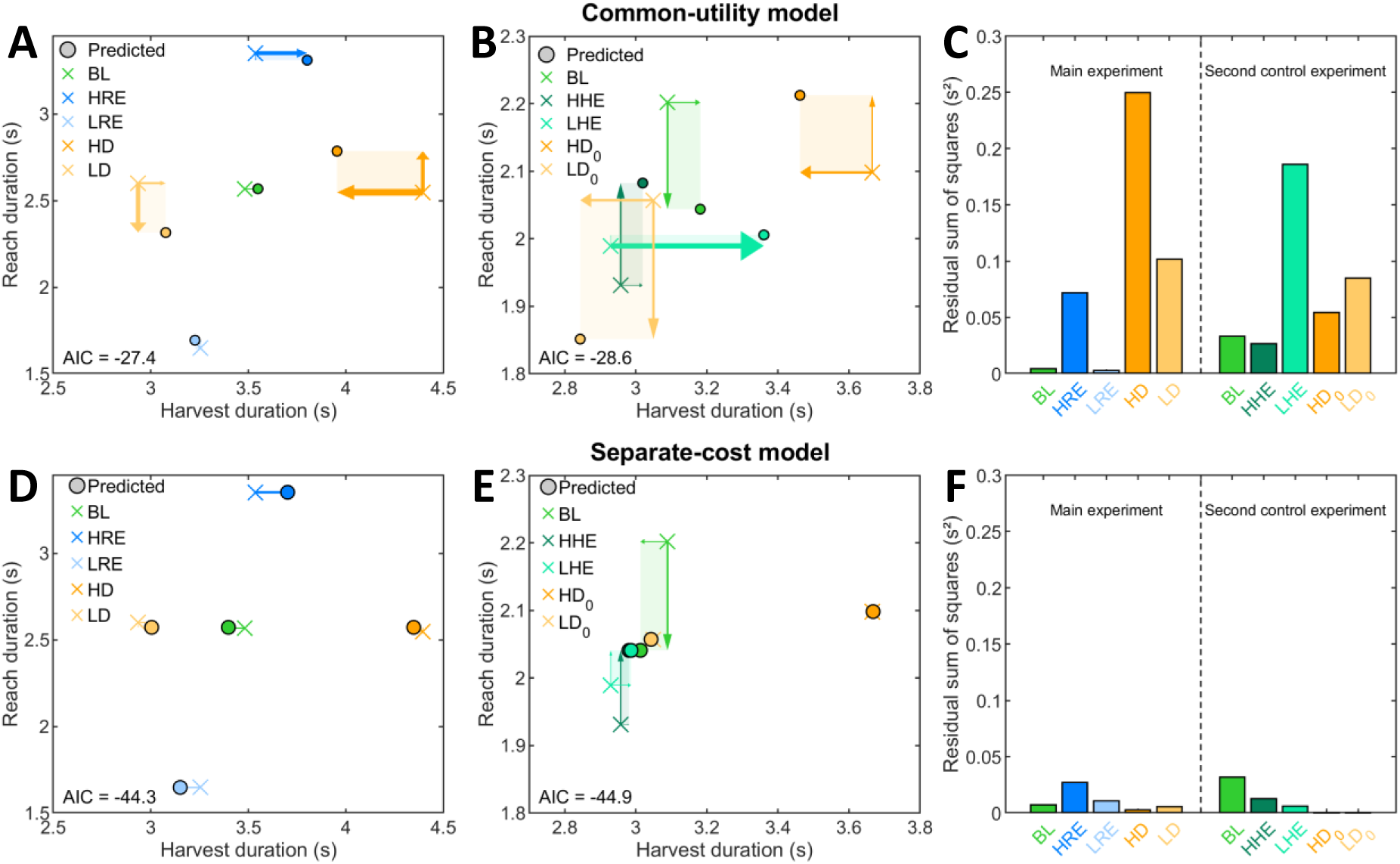
Model predictions of reach and harvest durations. (**A, B, D, E**) Colored crosses represent the median harvest (x axis) and reach (y axis) durations across all participants in the main (left panel) and second control (right panel) experiments. Predicted values from the common model (**A**, **B**) and the separate model (**D**, **E**) are shown as colored circles with black borders. Arrows and rectangles illustrate the deviation between model predictions and actual data. Residual sum of squares for each condition in the main experiment and second control experiment from the common (**C**) and separate (**F**) models. BL: Baseline (green); HRE: High Reach Effort (blue); LRE: Low Reach Effort (light blue); HHE: High Harvest Effort (dark green); LHE: Low Harvest Effort (light green); HD_0_: High Delay (orange); LD_0_: Low Delay (yellow).

**Table 1.**
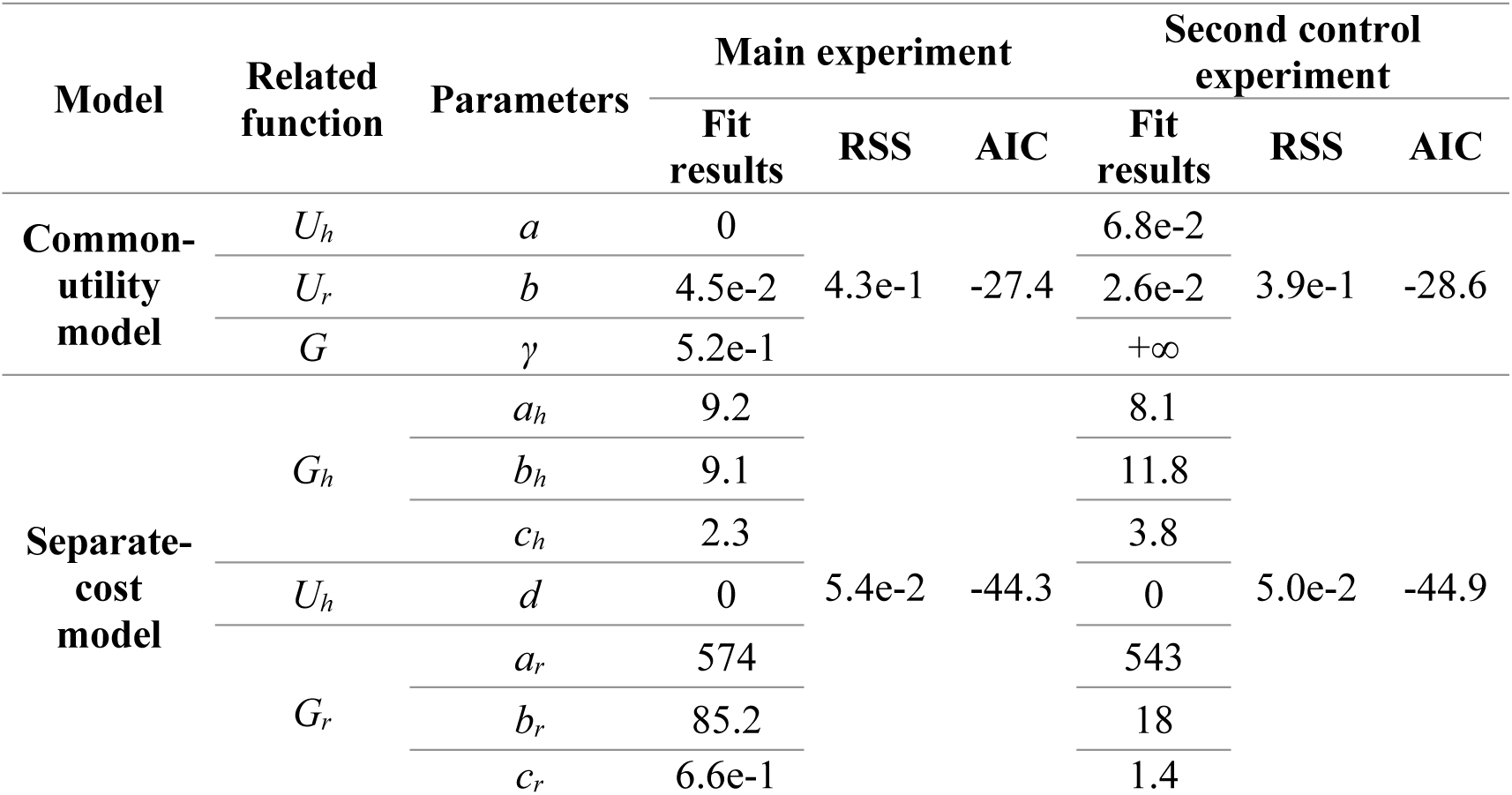
Overview of model fitting results. This table summarizes the best-fitting parameters, the *RSS* and the *AIC* for the common-utility and separate-cost models, in the main and second control experiments. The associated functions of each free parameter are also indicated. *U_h_*: harvest effort; *U_r_*: reach effort; *G*: temporal discount; *G_h_*: harvest’s cost of time; *G_r_*: reach’s cost of time.

For the common-utility model, the goodness of fit between the predicted data from the model and the observed data resulted in a total error of 657 milliseconds (*RSS* = 4.3e-1 s²; *AIC* = -27.4) in the main experiment (Fig. 6A). The best-fitting parameters were as follows: *a* = 0, *b* = 4.5e-2, and *γ* = 5.2e-1. Interestingly, *a* being 0 implies that the harvest effort *U_h_* was not taken into account to predict participants’ behavior. In the second control experiment (Fig. 6B), the goodness of fit between the predicted data from the model and the observed data resulted in a total error of 621 milliseconds (*RSS* = 3.9e-1 s²; *AIC* = -28.6). The best-fitting parameters were as follows: *a* = 6.8e-2, *b* = 2.6e-2, and *γ* → +∞. This γ value highlights that the model systematically improved by increasing the upper bound of the *γ* parameter. While it seems unrealistic, leading to very extreme temporal discounting, it still allows for the existence of a utility maximum, and thus, better optimal durations.

For the separate-cost model, the goodness of fit between the predicted data from the model and the observed data resulted in a total error of 231 milliseconds (*RSS* = 5.4e-2 s²; *AIC* = -44.3) in the main experiment (Fig. 6C). The best-fitting parameters were as follows: *a_h_* = 9.2, *b_h_* = 9.1, *c_h_* = 2.3, *d* = 0, *a_m_* = 574, *b_m_* = 85.2, and *c_m_* = 6.6e-1. It is noteworthy that *c_m_* < 1 implies that the time cost of reaching does not present a sigmoidal shape, but is rather hyperbolic. For the second control experiment (Fig. 6D), the goodness of fit between the predicted data from the model and the observed data resulted in a total error of 224 milliseconds (*RSS* = 5.0e-2 s²; *AIC* = -44.9). The best-fitting parameters were as follows: *a_h_* = 8.1, *b_h_* = 11.8, *c_h_* = 3.8, *d* = 0, *a_m_* = 543, *b_m_* = 18, and c*_m_* = 1.4. In both experiments, the scaling parameter *d* associated with *U_h_* systematically converged towards 0, suggesting that the harvest cost of effort was not critical to predict participants’ behavior. This may indicate that participants either did not perceive changes in harvest effort or did not consider them when adjusting their behavior. However, the former is unlikely, as additional analyses revealed that effort was objectively higher in the high harvest effort condition (conversely for the low harvest effort condition) (Fig. S4-5) and most participants subjectively noticed the difference (Table S1). Overall, these results suggest that the separate-cost model provided a better fit, even after taking into account its larger number of parameters compared to the common-utility model. A summary of the theoretical predictions and their comparisons with actual experimental observations is reported in Table 2. The best-fitting parameters were used to determine these theoretical predictions, giving the direction of changes whenever the effort or delays increase or decrease by an arbitrary amount (see Yoon et al., 2018 for a similar procedure, and Supplementary Video for a graphical illustration). It should be noted that indirect effects are not taken into account here. In the separate-cost model, for instance, an increase in reach effort is expected to increase reach duration, which, in turn, would affect harvest duration due to increased delays. Here, only the basic case was considered, in which this model predicts that a strict increase/decrease in reach effort with fixed reach duration should not affect harvest duration.

**Table 2.**
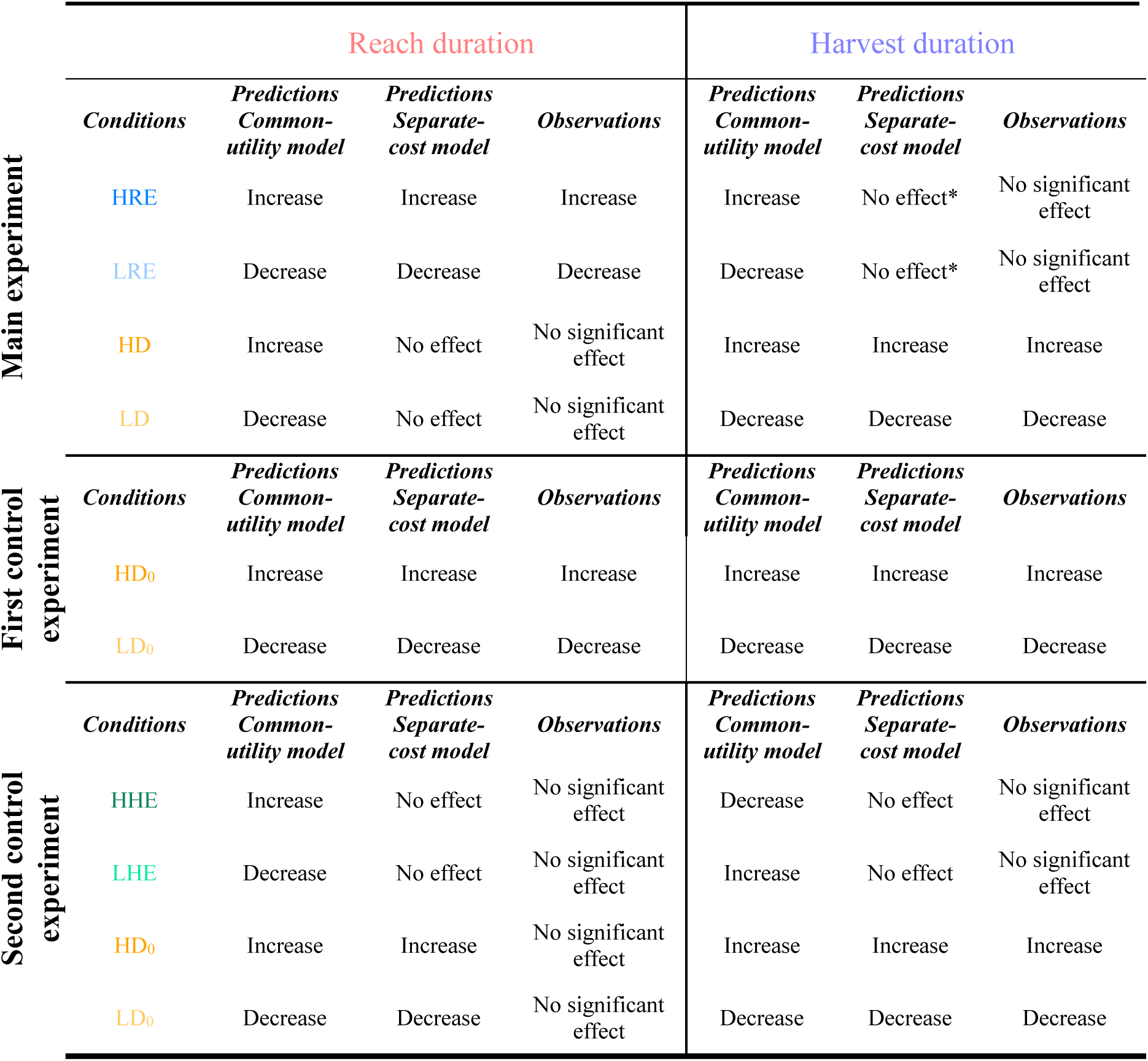
Summary of theoretical predictions and experimental observations. The direction of the predicted effects was determined from the parameters of the models fitted to the experimental data. *Indirect effects were not considered here; further details are provided in the main text. BL: Baseline; HRE: High Reach Effort; LRE: Low Reach Effort; HD: High Delay; LD: Low Delay; HHE: High Harvest Effort; LHE: Low Harvest Effort; HD_0_: High Delay; LD_0_: Low Delay.

To assess further whether the predictions of the common-utility and separate-cost models aligned with the experimental data, we also compared the observed and predicted changes in duration between the high- and low-effort conditions, and between the high- and low-delay conditions, separately for the harvest and reach phases. Specifically, we computed the difference in duration predicted by the common and separate models between the HRE-LRE and HD-LD conditions in the main experiment, and those predicted between the HHE-LHE and HD_0_-LD_0_ conditions in the second control experiment. These differences in predicted durations were then statistically tested using a one-sample Wilcoxon signed-rank test against the observed duration differences (Fig. 7).

**Figure 7.**
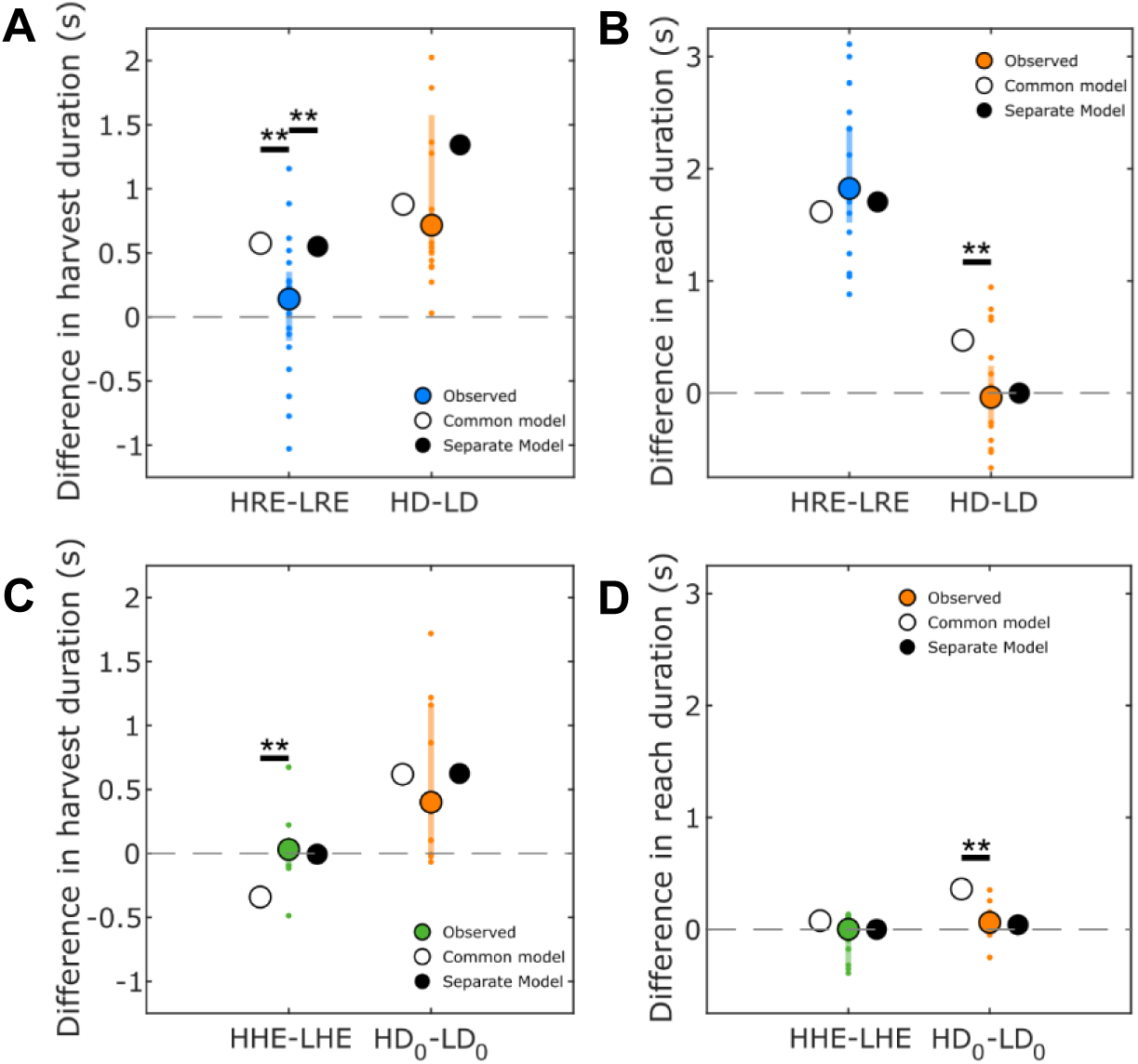
Experimental deviations from the predicted changes in duration. Difference in harvest (**A**) and reach (**B**) durations between the high- and low-reach effort conditions (HRE-LRE; in blue) and between the high- and low-delay conditions (HD-LD; in orange) in the main experiment. Difference in harvest (**C**) and reach (**D**) durations between the high- and low-harvest effort conditions (HHE-LHE; in green) and between the high- and low-delay conditions (HD_0_-LD_0_; in orange) in the second control experiment. These observed differences are compared with the predicted differences made by the common-utility model (white circle) and the separate-cost model (black circle). Colored circles represent median differences in duration, and bars represent interquartile ranges (n = 20 and n = 10 for each condition in the main and second control experiments, respectively). Individual data points are displayed as small colored dots. The stars represent p-values associated with the Wilcoxon signed-rank test. ** *p* < 0.01. HRE: High Reach Effort; LRE: Low Reach Effort; HHE: High Harvest Effort; LHE: Low Harvest Effort; HD_0_: High Delay; LD_0_: Low Delay.

Regarding harvest duration, in the main experiment (Fig. 7A), the observed HRE-LRE difference was significantly smaller than the difference predicted by both the common-utility model (observed: ΔT = 0.14 s, IQR = 0.54 s; predicted: ΔT = 0.58 s, W = 20, *p* = 0.002, r_rb_ = - 0.81) and the separate-cost model (predicted: ΔT = 0.55 s, W = 22, *p* = 0.002, r_rb_ = -0.79). In contrast, for the HD-LD comparison, the observed change in harvest duration did not significantly differ from the predictions of either the common-utility model (observed: ΔT = 0.72 s, IQR = 1.10 s; predicted: ΔT = 0.88 s, W = 111, *p* = 0.82, r_rb_ = 0.06) or the separate-cost model (predicted: ΔT = 1.34 s, W = 69, *p* = 0.18, r_rb_ = -0.34). In the second control experiment (Fig. 7C), the HHE-LHE difference was significantly different from the prediction of the common-utility model (observed: ΔT = 0.03 s, IQR = 0.20 s; predicted: ΔT = -0.34 s, W = 54, *p* = 0.004, r_rb_ = 0.96), but not from that of the separate-cost model (predicted: ΔT = -0.01 s, W = 33, *p* = 0.63, r_rb_ = 0.20). Finally, for the HD_0_-LD_0_ comparison, the observed change in harvest duration did not significantly differ from the predictions of either the common-utility model (observed: ΔT = 0.40 s, IQR = 1.17 s; predicted: ΔT = 0.62 s, W = 23, *p* = 0.70, r_rb_ = -0.16) or the separate-cost model (predicted: ΔT = 0.62 s, W = 23, *p* = 0.70, r_rb_ = -0.16).

Regarding reach duration, in the main experiment (Fig. 7B), the observed HRE-LRE difference did not significantly differ from the predictions of either the common-utility model (observed: ΔT = 1.82 s, IQR = 0.84 s; predicted: ΔT = 1.62 s, W = 154, *p* = 0.067, r_rb_ = 0.47) or the separate-cost model (predicted: ΔT = 1.70 s, W = 140, *p* = 0.19, r_rb_ = 0.33). For the HD-LD comparison, however, the observed change in reach duration was significantly different from the prediction of the common-utility model (observed: ΔT = -0.04 s, IQR = 0.52 s; predicted: ΔT = 0.47 s, W = 17, *p* = 0.001, r_rb_ = -0.84), but not from that of the separate-cost model (predicted: ΔT = 0 s, W = 99, *p* = 0.82, r_rb_ = -0.06). In the second control experiment (Fig. 7D), the observed HHE-LHE difference did not significantly differ from the predictions of either the common-utility model (observed: ΔT = 0 s, IQR = 0.43 s; predicted: ΔT = 0.08 s, W = 15, *p* = 0.23, r_rb_ = -0.45) or the separate-cost model (predicted: ΔT = 0 s, W = 20, *p* = 0.49, r_rb_ = -0.27). Finally, for the HD_0_-LD_0_ comparison, the observed change in reach duration was significantly different from the prediction of the common-utility model (observed: ΔT = 0.06 s, IQR = 0.20 s; predicted: ΔT = 0.36 s, W = 0, *p* = 0.002, r_rb_ = -1), but not from that of the separate-cost model (predicted: ΔT = 0.04 s, W = 30, *p* = 0.85, r_rb_ = 0.09).

In sum, these analyses indicate that the common-utility model tended to overestimate the changes in duration induced by time (HD-LD; HD_0_-LD_0_) and effort (HRE-LHE; HHE-LHE) modulations. By contrast, the separate-cost model better accounted for the observed pattern of effects, although both models failed to capture the relative stability of harvest duration following reach-effort modulations (Fig. 7A)

Interestingly, the sigmoidal shape found for the cost of time of the harvest and reach phases (only for the second control experiment for the latter), allows us to compute the time *t_i_* at which the inflection point happens (see methods). This moment marks the switch between a fast exponential increasing cost of time, toward a more hyperbolic cost of time. Since all conditions in an experiment share the same adjusted parameters, the inflection point is also shared and represents somehow an average between all conditions. For the main experiment, only the time cost of harvesting presents a sigmoidal shape, with an inflection point located at 6.0 seconds. Of note, across conditions, the average reach’s time offset *o_r_* is 0 (as no delay occurs before the reach phase), and the harvest’s time offset *o_h_* is 5.5 seconds. For the second control experiment, the time cost of reaching shows an inflection point at 4.3 seconds and the time cost of harvesting inflection point is located at 10.2 seconds. Across conditions, the average reach’s time offset *o_r_* is 3 seconds, and the harvest’s time offset *o_h_* is 8.1 seconds. These findings point to several interesting observations (Fig. 8). First, the inflection point location increases linearly with the time offset (Fig. 8A-B). Second, the inflection point aligns closely with the beginning of the corresponding phase, although with a certain tendency to appear during the phase, especially with large average time offsets (Fig. 8A). Consequently, time offsets are primarily captured by an exponential cost of time (before the inflection point), while a hyperbolic cost of time more accurately characterizes the value of the time elapsed during actions (Fig. 8B). The latter also highlights why the time cost of reaching is only hyperbolic in the main experiment, as no actual delay (and therefore no offset) occurs before. Altogether, delays occurring before movement and decision play an important role in their behavioral invigoration, as they increase exponentially the cost of time.

**Figure 8.**
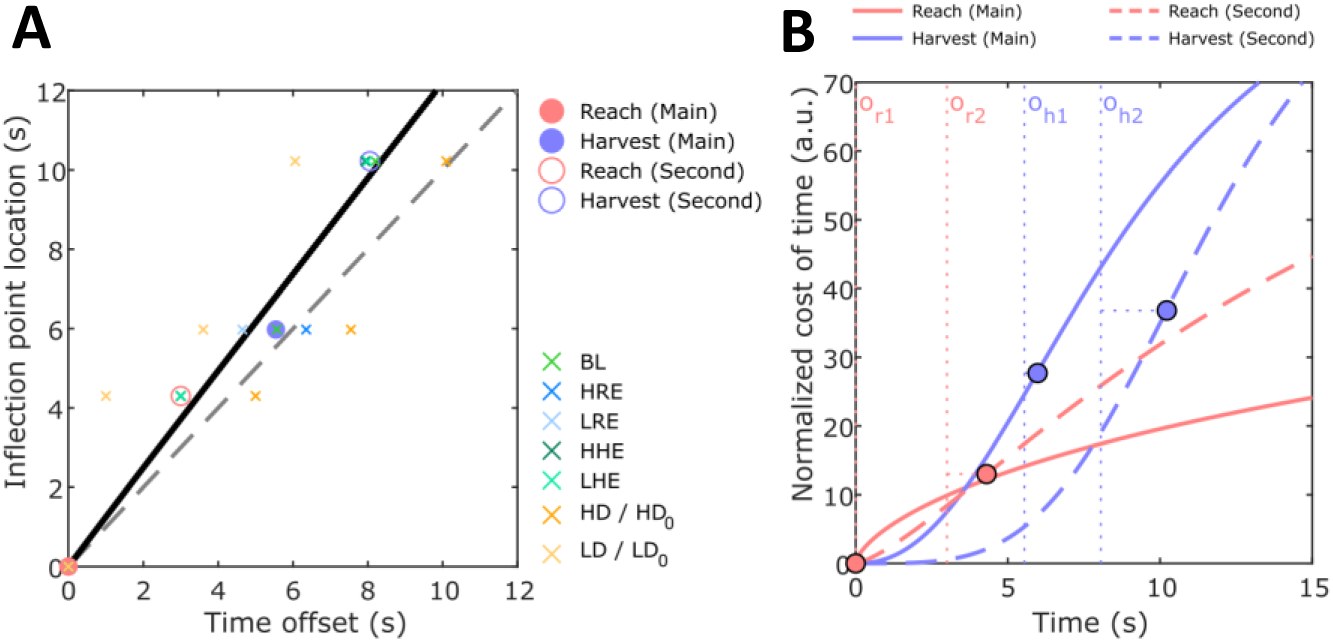
Inflection points of the costs of time according to the time offset. **(A)** Large colored circles represent the inflection point of each cost of time in relation to the time offset (red for reach phases, blue for harvest phases) for the main experiment (filled circles) and the second control experiment (empty circles). Small colored crosses represent each condition’s time offset with the corresponding inflection point. A power-law function was fitted (black line) such that *t_i_*(*o*) = *a*_1_*o^a^*^2^, where o is the average time offset and *a_1_* and *a_2_* are two free parameters. Fit result: R² = 0.98, RSS = 1.18 s²; *a_1_* = 1.26; *a_2_* = 0.99. The dotted line represents theoretical values of the inflection point if it occurs exactly at the start of the phase of interest. (**B**) Representation of each cost of time of the reach (red) and harvest (blue) phases found for the main (solid lines) and second control experiment (dashed lines). The costs of time were normalized by their parameter *a*. Dotted lines represents the corresponding average time offsets of the reach (*o_r_*_1_, *o_r_*_2_ respectively for the main and second control experiment) and harvest phases (*o*_*h*1_, *o*_*h*2_ respectively for the main and second control experiment). Note how the inflection point location (colored circles) and the steepness of the curve increase with the time offset. BL: Baseline (green); HRE: High Reach Effort (blue); LRE: Low Reach Effort (light blue); HD / HD0: High Delay (orange); LD / LD0: Low Delay (yellow); HHE: High Harvest Effort (dark green); LHE: Low Harvest Effort (light green).

## Discussion

In the current study, we investigated whether the vigor of movement and decision-making are governed by a common or a separate control mechanism in a reward-oriented task designed to mimic human foraging behavior. In our paradigm, the reach phase indexes movement vigor, whereas the harvest phase captures decision vigor, reflecting how long participants choose to exploit a rewarding patch before relocating (Sukumar et al., 2024; Yoon et al., 2018). Behavioral analyses and modelling were conducted to explore how time and effort variations in the reach and harvest phases affect their respective durations. At the behavioral level, we found that reach and harvest durations were affected by preceding delays in the current trial (although only slightly for the reaching as it was not replicated in the second control experiment). However, reach duration was not affected by delays occurring *after* the movement within a trial. In addition, reach effort modulated the duration of the reach phase but did not change harvest duration. Unexpectedly, no change in duration for either the reach or harvest phases was observed after changes in harvest effort. At the inter-individual level, we found a strong consistency between reach durations across conditions, as well as for harvest durations. However, no relationship was found between individual’s reach duration and harvest duration, thereby showing that vigorous participants during movement were not necessarily the most vigorous during decision-making. Although the common-utility model performed relatively well overall, it failed to account for some observations. The alternative model, assuming that movement and decision stem from separate, yet inter-dependent, optimization processes, provided a better account of the experimental findings in this specific environmental context by allowing a more flexible regulation of their vigor (Table 2). Below we discuss the implications and limitations of our study.

### Effect of reach and harvest efforts

Consistent with previous findings, increasing reaching effort led to longer reach durations (Bruening et al., 2024; Sukumar et al., 2024). However, contrary to the predictions of the common-utility model tested here (based on the global capture rate maximization), reaching effort did not significantly affect harvest duration. This aligns with (Sukumar et al., 2024), who found that changes in the environment’s movement effort did not alter the amount of reward harvested. In their study, discrete reward paired with salient feedback may have encouraged participants to collect a fixed number of rewards. Here, the use of a continuous, feedback-free reward suggests that this lack of effect rather reflects individuals’ decision-making behavior, decoupling reaching effort and harvest duration, than an unintended consequence of reward-related feedback. In support of this interpretation, recent work revealed that lesions of the dorsal striatum in rats led to longer movement durations by enhancing sensitivity to effort, without affecting decision duration (Morvan et al., 2024).

Strikingly, variations in harvesting effort affected neither movement nor decision durations in our task. Similar deviations from the common-utility model predictions were reported by (Yoon et al., 2018), who found that after a history of high effort induced by gazing at highly eccentric images, participants increased both harvest duration (in agreement with the model) and saccade speed (in disagreement with the model). This effect was interpreted by the justification of effort, whereby higher effort heightens the subjective value of the reward (Clement et al., 2000), thereby reducing movement duration (Summerside et al., 2018). A similar compensation in our study could then explain why increasing harvest effort within a block did not slow the reaching. More unexpectedly, harvesting effort also failed to affect the duration of the harvest phase itself. Since shorter harvests directly reduce reward, participants may have been reluctant to adjust this phase. Consistent with this, the subjective value of reward is seemingly discounted similarly by moderate isometric efforts (10-50% of the MVC) (Burke et al., 2013; Hartmann et al., 2013). Hence, a possible hypothesis would be that the perceived effort of harvest was outweighed by the reward value. For instance, Morvan et al. (2024) showed in rats that decision duration depends more on reward and satiety than on effort or fatigue. The effort induced by high eccentricity in the study of (Yoon et al., 2018) may have been of greater magnitude than the effort variation implemented in our experiment. Rather than attributing this outcome to the justification of effort, we speculate that the harvesting effort applied here may have been insufficient to disrupt participants’ behavior, and that much greater efforts could have shortened harvest duration at some point. Although clear changes were observed in the effort exerted during harvest (Fig. S4-5), it was not substantial enough to influence participants’ harvesting behavior.

Altogether, it appears that there are still uncertainties regarding the integration of effort in the regulation of movement and decision-making vigor. Here, movement effort seems to carry more weight than decision effort, although they were of very similar physical magnitudes in our task. This suggests that reward may modulate or even override the behavioral changes typically associated with increased effort, specifically regarding decision duration, which was directly linked to reward collection.

### The behavioral value of time

While the common-utility model predicts similar effects of delays on harvest and reach durations, we found respectively marked and small effects. Importantly, this observation was true for the latter only if the delays appeared before the reach phase. This phenomenon is well captured by considering a time-growing signal—termed the cost of time—that urges participants to increase their vigor in each phase; its onset coincides with the beginning of each trial in our task. Interestingly, the sigmoidal cost-of-time function identified in prior works exhibited an inflection point located near the onset of each phase, with a time shift reflecting the elapsed duration between the start of the trial and the beginning of that phase. Consequently, the cost of time used to determine behavioral invigoration was mostly hyperbolic, whereas it tended to increase exponentially during the preceding delay periods. This cost-of-time function underlying behavioral invigoration may reflect a time-sensitive “boredom” (Berret et al., 2018) or “urgency” signal (Thura et al., 2014), remaining relatively flat initially and increasing quickly with time. When it serves to determine reach and harvest durations, the cost of time would have therefore undergone an increase or been magnified, resulting in more vigorous actions. The cost of time implicitly represents the temporal discount of the reward (Shadmehr, 2010; Summerside et al., 2018). As such, an increase in the available reward to be harvested could induce steeper time-costs, thereby affecting both reach and harvest durations regardless of their sequential order (Yoon et al., 2018; Sukumar et al., 2024). Reward manipulation may therefore inherently induce co-regulation by directly shaping the behavioral value of time (i.e., the cost of time).

Remarkably, according to this cost of time hypothesis, the separate-cost model implies nevertheless an inter-dependent regulation of reach and harvest durations also outside reward manipulation. Indeed, harvest duration should have been affected by reaching effort, since greater effort leads to longer reach durations and thus delays the harvest onset. Yet, we observed no such effect. This absence could stem from our biased perception of time during movement, which tends to be compressed by action (Tomassini et al., 2014). Increasing reaching effort also reduce the velocity of the visual stimulus (i.e., cursor position), potentially shortening perceived duration (Brown, 1995; Tomassini et al., 2011) and counterbalancing the actual changes in duration. Constantino and Daw (2015) showed that longer travel delays prolong subsequent harvests, but their paradigm involved effortless, simulated movements without actual action. Hence, in our experiment, despite changes in reach duration in the high/low reaching effort conditions, participants appeared relatively insensitive to these variations, while remaining highly sensitive to delays experienced during inactivity. Since the cost of time necessarily involves subjective time perception, these phenomena could explain this discrepancy.

Altogether, our results provide indirect evidence for the existence of a cost of time that is sensitive to delays recently experienced by the participants in the task (or trial). Movement and decision invigoration would then result from distinct optimization processes balancing a shifted cost of time with other task-relevant costs (e.g., effort, reward, accuracy etc.). It is worth mentioning that while these optimizations are mathematically separate, they can nevertheless influence one another due to an inherent integration of past temporal delays within the current cost of time. The time shift induced by previously experienced delays may still cause some degree of inter-dependence between movement and decision vigor.

### Flexible invigoration of movement and decision

It has been proposed that the co-regulation between decision and movement vigor stems from an urgency signal (Thura et al., 2014), initially described in the context of decision-making (Cisek et al., 2009). This urgency signal is described as an evidence-independent signal that grows over time during deliberation, pushing the action-related neural activity closer to the decision threshold, thereby speeding choice commitment, especially when sensory evidence is weak. Previous findings showed that this urgency signal jointly invigorates both decision and movement to optimize the capture rate (Fievez et al., 2024; Thura et al., 2014; Thura, 2020). However, decoupling may occur when the default co-regulation would be detrimental to the utility maximization (see (Thura et al., 2025) for a review). For example, a compensatory trade-off between decision and movement duration can be elicited when slowing down the movement to maintain accuracy, while deciding quickly is required to maximize a pure reward rate (Reynaud et al., 2020). Hence, the urgency signal is not a rigid drive and can be flexibly gated by higher-level task goals when co-regulation becomes counterproductive. Our task environment may have favored a decoupling of movement and decision vigor. Specifically, the fixed time constraint within each block may have provided a strong incentive to override the default invigoration driven by the urgency signal, thereby promoting more independent optimization of reach and harvest durations. In tasks where decision and movement share the same urgency signal, however, our model based on a shifted cost of time could actually explain co-regulation. For instance, reaction time preceding movement onset can be considered as a decision-making process, thereby inducing a short delay before task execution. The presence of a steep time-cost signal that rapidly reaches the threshold would yield both short reaction times and short movement times, consistent with any experimentally-observed co-regulation (Takikawa et al., 2002; Haith et al., 2012; Reppert et al., 2018). Additional analyses of reach reaction time corroborate this hypothesis, as we observe changes in the same direction as reach duration after time and effort manipulations (Fig. S2). Conversely, when movement and decision phases are tightly constrained or segregated (i.e., governed by distinct costs of time), the brain may optimize their durations independently. Thura et al. (2025) recently proposed an interesting view of the neurophysiological correlates allowing such flexibility between co-regulation and decoupling: the recruitment of inhibitory control mechanisms, involving the hyper-direct pathway connecting the frontal cortex to the subthalamic nucleus, can exert a large suppressive influence on motor activity. This entails an increase in the decision threshold, allowing deliberation to proceed more slowly and accurately, even when the motor system is prepared for vigorous movement.

The utility-framework suggests, in contrast, that movement and decision are strongly and rigidly inter-dependent. Thus, it was recently hypothesized that an individual exhibiting high movement vigor should also have high decision vigor (Reppert, 2025). While our findings confirm consistent individual movement vigor (Labaune et al., 2020) and highlight the presence of decision vigor, we found little correlation between the two, suggesting distinct underlying invigorations. Previous work has shown that vigor may not represent a universal individual trait by reporting independent vigor for arm reaching and eye saccades at the inter- and intra-individual levels (Labaune et al., 2020; Thura, 2020). This raises the question of whether the co-regulation between decision and movement vigor, observed in eye-saccade paradigms for instance, generalizes to other types of movement. It is noteworthy that the lack of inter-individual consistency may have been fostered by an environment that is not conducive of reach and harvest co-regulation. Future studies should assess whether a co-regulation between movement and decision vigor emerging at the intra-individual level can also be observed at the inter-individual level. To support this hypothesis, supplementary analyses show that the apparent co-regulation of reaction time and reach duration found at the intra-individual level is indeed also reflected at the inter-individual level (Fig. S2-3).

In conclusion, we investigated movement and decision invigoration in a foraging-like task balancing time and effort. We tested two alternative models to explain the behavioral vigor of participants. Our findings revealed a decoupling between reach and harvest durations in this task, which was better predicted in this controlled environmental context by a model allowing separate yet inter-dependent optimizations of decision and movement vigor than by a common-utility model. This suggests that the common-utility model, while performing overall well, may need to be refined to allow for greater flexibility in movement and decision invigoration. Behavioral vigor may thus rely on a mechanism involving a cost of time that is sensitive to recently experienced delays, thereby indirectly linking movement and decision-making. The emergence of decoupling or co-regulation would depend on whether shared or independent time-cost signals govern movement and decision invigoration.

## Data availability

The data underlying the graphs presented in the figures of the main manuscript and in Supplementary Information are available as Supplementary Data upon publication. Additional data that support the results of this study are available from the corresponding author upon reasonable request and under a formal data-sharing agreement.

## Code availability

The codes used in this study are available from the authors upon reasonable request.

## Acknowledgement

This work was supported by the French National Agency for Research (grant ANR-22-CE37-0010, BasalCost project).

## Author contributions

Author contributions: A.C., A.B. and B.B. conceived the experiment; A.C. collected the data; A.C. and B.B. analyzed the data and discussed the results; A.C. and B.B. wrote the manuscript; All authors revised the manuscript.

