## Supplementary information for "Evidence for flexible regulation of movement and decision vigor in a reward-oriented task"

#### Time-related supplementary analyses

To further assess the effect of time and effort changes on participants' behavior, trial-by-trial presentation of reach and harvest durations are provided (Figure S1), as well as additional analyses on reach reaction times (RT) (Figure S2). To compute RT, the raw positions were smoothened using a zero-phase 5th-order Butterworth low-pass filter with 3 Hz cutoff frequency. Velocity was then computed via numerical differentiation. It is noteworthy that in the main experiment the reach phase is preceded by the harvest phase, with no marked stop between them, which can lead to situations where the speed does not decrease substantially. Hence, reaction time was defined as the duration between the reach phase onset and the last point under 5 cm/s before the peak velocity. This conservative approach allowed us to only discard four trials from these analyses across participants in the main experiment. Also, given the highly cyclic nature of the experimental paradigm, anticipation of target appearance is expected from participants, specifically in the control experiments where reach phases are preceded by a delay. Hence, movement onset was determined using a time window extending up to 500 ms before the start of the reach phase, allowing possible negative values for reaction times (relative to reach phase onset). RTs were then compared between conditions for each experiment using Wilcoxon signed-rank tests.

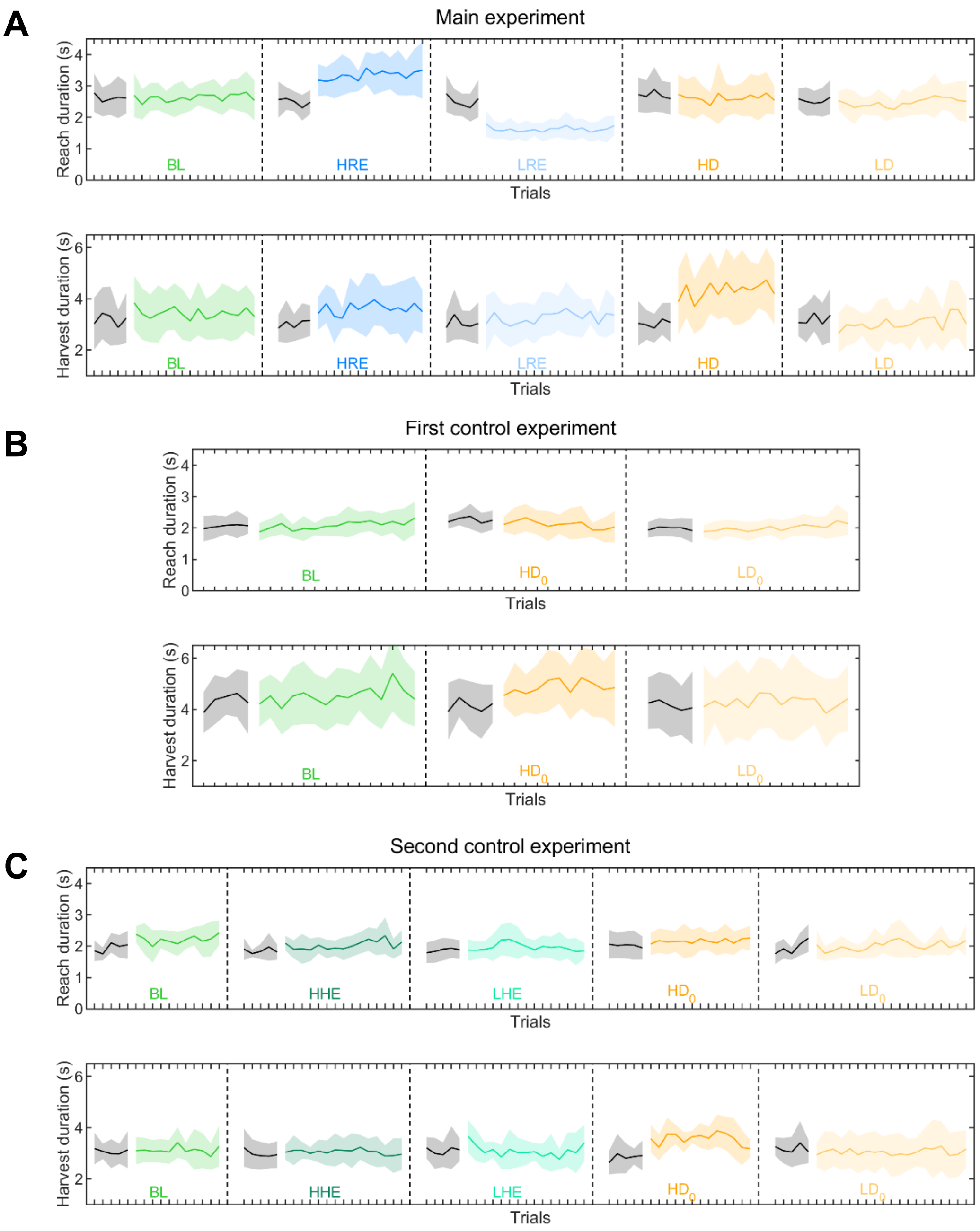


**Figure S1. Trial-by-trial reach and harvest durations.** Reach and harvest durations for each trial and condition of the main (**A**), first control (**B**), and second control (**C**) experiments. Thick lines represent median durations (n = 20, n = 14, n = 10 respectively for the main, first control and second control experiments) and the lighter areas surrounding the medians represent the median absolute deviations. Black lines represent wash-out trials (before the introduction of time or effort manipulation). Since participants did not perform the same number of trials within a block, only the duration of the trials performed by all participants was taken into account. BL: Baseline; HRE: High Reach Effort; LRE: Low Reach Effort; HD: High Delay; LD: Low Delay; HHE: High Harvest Effort; LHE: Low Harvest Effort; $HD_0$: High Delay; $LD_0$: Low Delay.

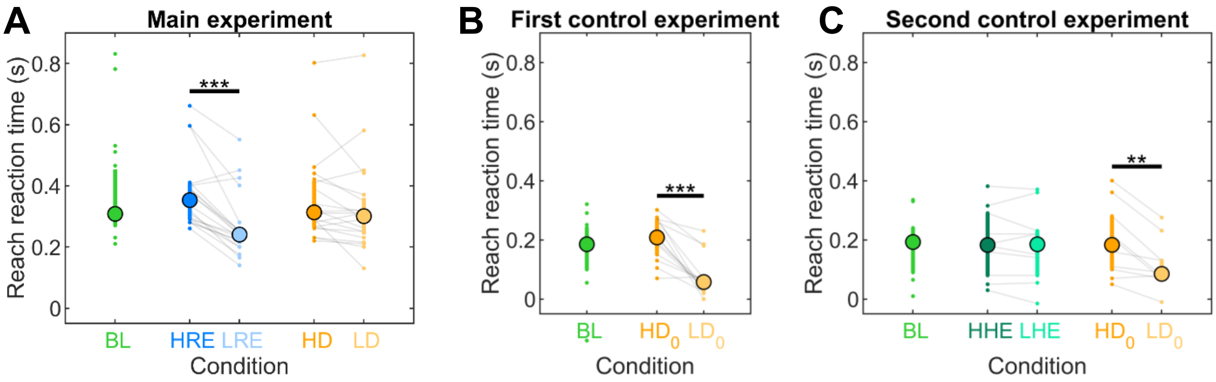


**Figure S2. Effects of time and effort changes on reach reaction times.** Reach reaction time for each condition of the main (**A**), first control (**B**), second control (**C**) experiments. Colored circles represent median durations, and bars represent interquartile ranges (n = 20, n = 14, n = 10 respectively for the main, first control and second control experiments). Individual data points are displayed as small colored dots. Thin grey lines connect compared observations across conditions. The stars represent p-values associated with Wilcoxon signed-rank tests. * $p < 0.05$, ** $p < 0.01$, *** $p < 0.001$. BL: Baseline; HRE: High Reach Effort; LRE: Low Reach Effort; HD: High Delay; LD: Low Delay; HHE: High Harvest Effort; LHE: Low Harvest Effort; $HD_0$: High Delay; $LD_0$: Low Delay.

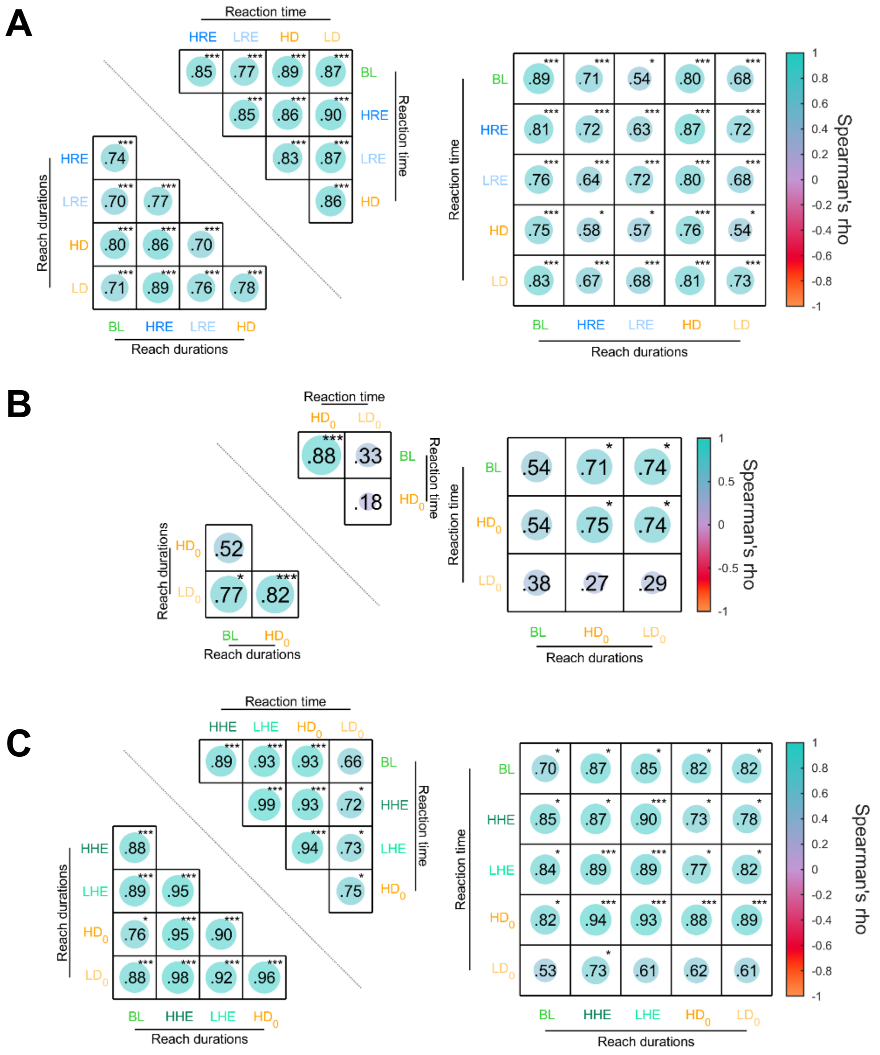


**Figure S3. Inter-individual consistencies of reach duration and reaction time.** Relationships between individuals' reach duration and reaction time for all pairs of conditions of the (**A**) main, (**B**) first control and (**C**) second control experiments. Left panel: heatmap of the correlations separately for reach duration (lower part) and reaction time (upper part) between each pair of conditions. Right panel: heatmap of the correlations between reach duration and reaction time for each pair of conditions. Spearman correlation coefficients (rho) are reported for each correlation. The stars represent p-values associated after correction for multiple comparisons using the Benjamini-Hochberg procedure (Ntests = 45 for the main and second control experiments, and Ntests = 15 for the first control experiment).* $p < 0.05$, ** $p < 0.01$, *** $p < 0.001$. BL: Baseline; HRE: High Reach Effort; LRE: Low Reach Effort; HHE: High Harvest Effort; LHE: Low Harvest Effort; $HD_0$: High Delay; $LD_0$: Low Delay

#### Effort-related supplementary analyses

Analyses of the extensor carpi radialis EMG signal (Figure S3) were performed to assess whether the changes in terms of effort were significant between conditions. To account for individual variations, the EMG signal was normalized by the highest absolute EMG value obtained during three maximal voluntary contraction against the robotic exoskeleton. To get an estimate of the cumulative effort exerted throughout the duration of reach and harvest phases, the EMG-time integral of the normalized signal was computed for each phase. The median of the resulting EMG-time integral was computed for each condition and then compared between conditions for each experiment using Wilcoxon signed-rank tests. Complementary analyses on the torque-time integral computed from the applied torque during each phase are also provided to provide a different estimate of the effort exerted during reach and harvest phases (Figure S4).

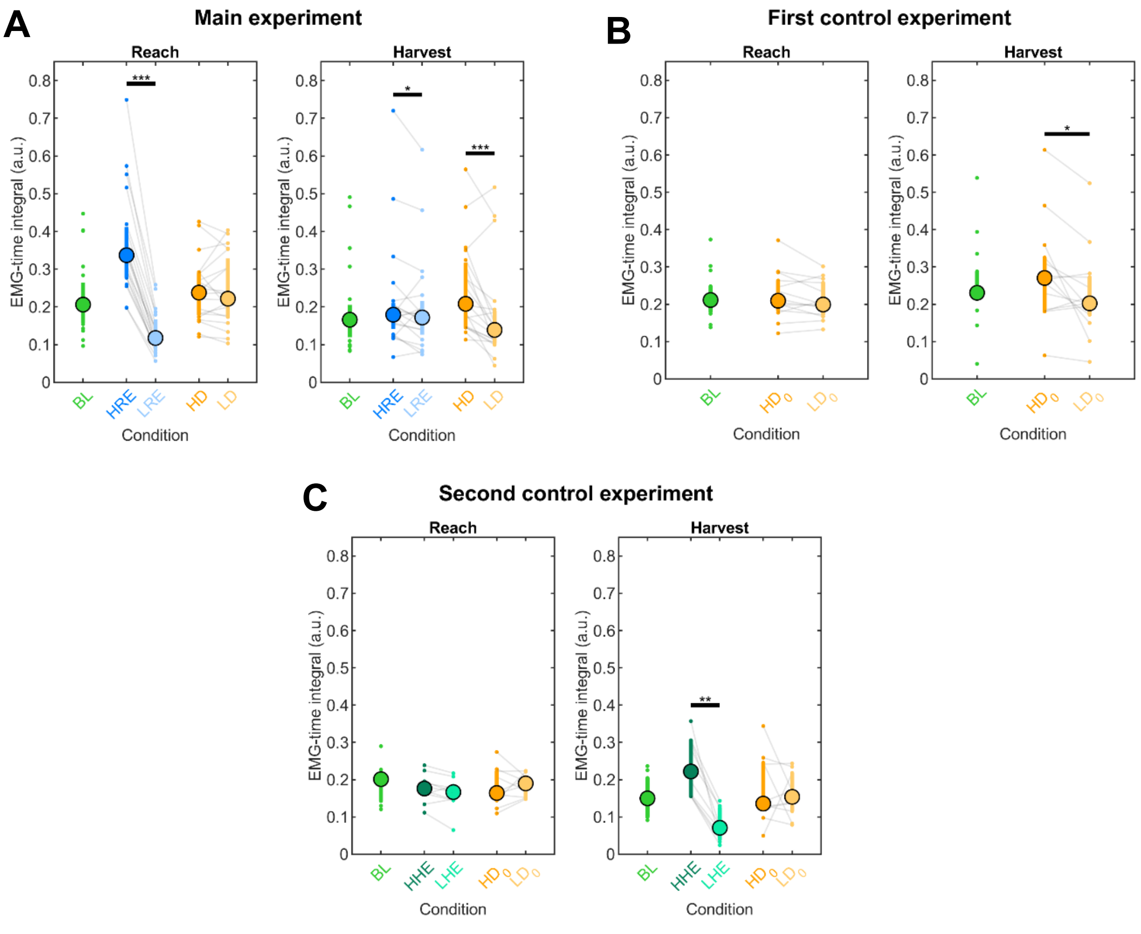


**Figure S4. Estimation of the effort exerted during reach and harvest phases from the EMG.** EMG-time integral for each condition of the main (**A**), first control (**B**) and second control (**C**) experiments. Colored circles represent median EMG-time integral values, and bars represent interquartile ranges (n = 20, n = 14, n = 10 respectively for the main, first control and second control experiments). Individual data points are displayed as small colored dots. Thin grey lines connect compared observations across conditions. The stars represent p-values associated with Wilcoxon signed-rank tests. * $p < 0.05$, ** $p < 0.01$, *** $p < 0.001$. BL: Baseline; HRE: High Reach Effort; LRE: Low Reach Effort; HD: High Delay; LD: Low Delay; HHE: High Harvest Effort; LHE: Low Harvest Effort; $HD_0$: High Delay; $LD_0$: Low Delay.

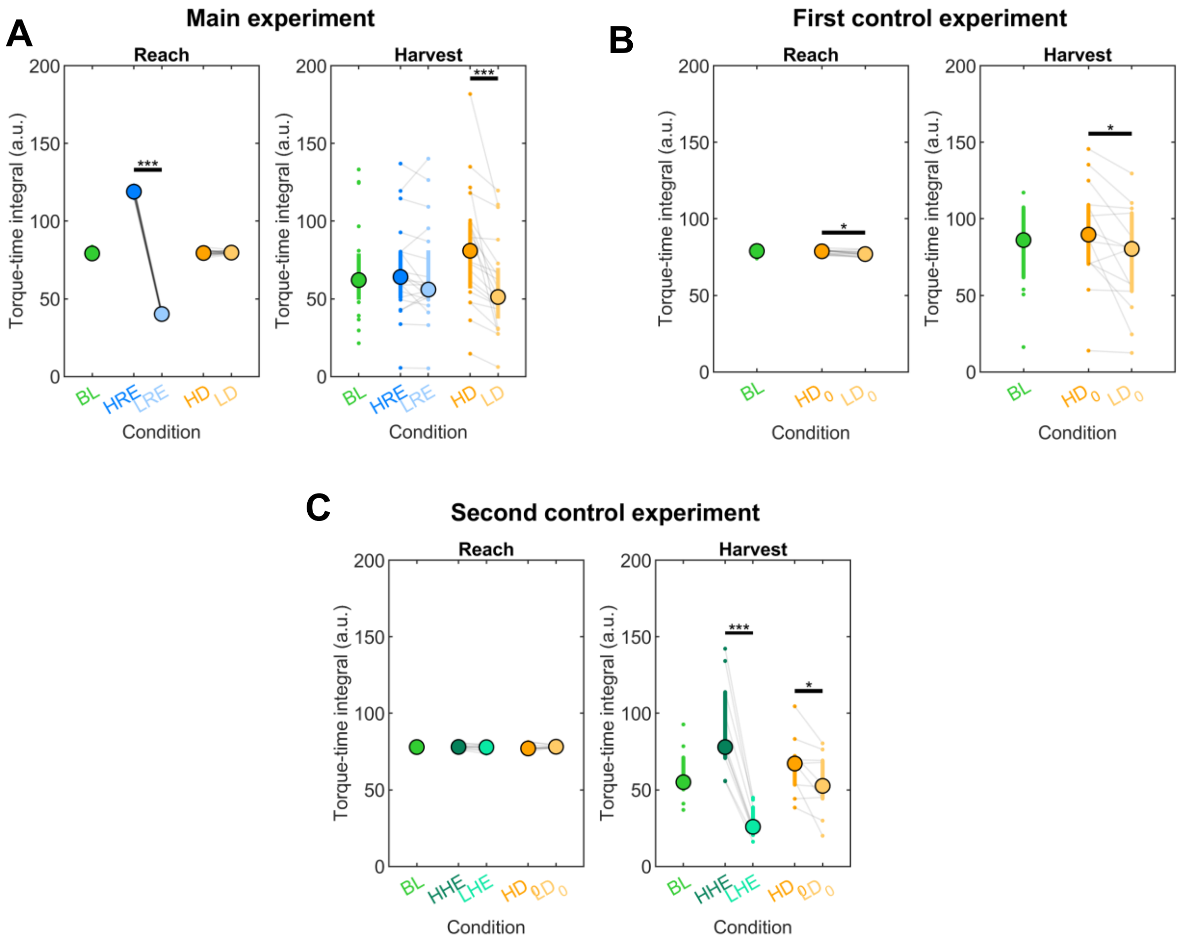


**Figure S5. Estimation of the effort exerted during reach and harvest phases from the applied torque.** Torque-time integral for each condition of the main (**A**), first control (**B**) and second control (**C**) experiments. Colored circles represent median torque-time integral values, and bars represent interquartile ranges (n = 20, n = 14, n = 10 respectively for the main, first control and second control experiments). Individual data points are displayed as small colored dots. Thin grey lines connect compared observations across conditions. Note that the theoretical value of the torque-time integral of the reach phase is constant for each condition (see the modeling section of the main manuscript), which accounts for the low interindividual variability compared to the harvest. The stars represent p-values associated with Wilcoxon signed-rank tests. * $p < 0.05$, ** $p < 0.01$, *** $p < 0.001$. BL: Baseline; HRE: High Reach Effort; LRE: Low Reach Effort; HD: High Delay; LD: Low Delay; HHE: High Harvest Effort; LHE: Low Harvest Effort; $HD_0$: High Delay; $LD_0$: Low Delay.

#### Subjective assessment

| *Experiments* | *Conditions* | | | |
|---|---|---|---|---|
| **Main experiment** | HRE | LRE | HD | LD |
| Number of aware participants (/20) | 16 | 15 | 17 | 18 |
| **First control experiment** | | | $HD_0$ | $LD_0$ |
| Number of aware participants (/14) | | | 14 | 13 |
| **Second control experiment** | HHE | LHE | $HD_0$ | $LD_0$ |
| Number of aware participants (/10) | 9 | 8 | 10 | 9 |

**Table S1. Explicit awareness of time and effort modifications.** Number of participants that correctly reported being aware of effort or delay changes compared to baseline and wash-out trials.

### Details about the models

In this section, a representation of how the various costs involved in the separate-cost model are organized within a typical trial of the second control experiment, showing how delays and reach phase duration can influence harvest duration through temporal offsets (Figure S6). Then, details about the supplementary video are provided, which illustrates how both models can lead to co-regulation or decoupling of reach and harvest durations. Finally, linear mixed model analyses were also performed (*lme4* package in R) to assess if the average torque is a good predictor of the maximum torque (Figure S7). We computed the median (across all trials for each condition) of the average normalized (by $\tau_{MVC}$) torque $\overline{\mu}$ and of the maximum normalized torque $\mu_{max}$ for each condition and participant. The median of $\mu_{max}$ was set as the dependent variable, with fixed and random effects for both the median of $\overline{\mu}$ and intercept, grouped by condition. The resulting equation is: $\mu_{max}$ ~ 1+ $\overline{\mu}$+(1+$\overline{\mu}|Condition$). The restricted maximum likelihood estimation (REML) was used to estimate the parameters.

*For the main experiment*, the fixed effect coefficient of intercept (Estimate = 3.65, SE = 1.53, df = 44.1, t = 2.39, $p = 0.022$) contributed significantly to predicting $\mu_{max}$, as well as the fixed effect coefficient of $\overline{\mu}$ (Estimate = 1.41, SE = 0.06, df = 4.88, t = 23.2, $p < 0.001$), giving a marginal $R^2$ of 0.86. The contribution of random effects for intercept and $\overline{\mu}$ was relatively poor, explaining only 4.8% additional variance (conditional $R^2 = 0.91$).

*For the first control experiment*, the fixed effect coefficient of intercept did not contribute significantly to predicting $\mu_{max}$ (Estimate = 3.24, SE = 2.31, df = 15.4, t = 1.40, $p = 0.18$), contrary to the fixed effect coefficient of $\overline{\mu}$ (Estimate = 1.29, SE = 0.06, df = 6.55, t = 20.44, $p < 0.001$), giving a marginal $R^2$ of 0.92. The contribution of random effects for intercept and $\overline{\mu}$ was negligible and did not explain additional variance.

*For the second control experiment*, the fixed effect coefficient of the intercept did not contribute significantly to predicting $\mu_{max}$ (Estimate = 2.65, SE = 1.63, df = 21.6, t = 1.62, $p$ = 0.12), contrary to the fixed effect coefficient of $\overline{\mu}$ (Estimate = 1.32, SE = 0.04, df = 10.2, t = 30.1, $p < 0.001$), giving a marginal $R^2$ of 0.95. The contribution of the random effects for intercept and $\overline{\mu}$ was negligible, explaining only 0.2% additional variance.

Overall, the model fitted well for all experiments. Minor contributions of random effects grouped by condition suggest that the median of the average normalized torque $\overline{\mu}$ is a good predictor of the median of the maximal normalized torque $\mu_{max}$, regardless of condition and experiment.

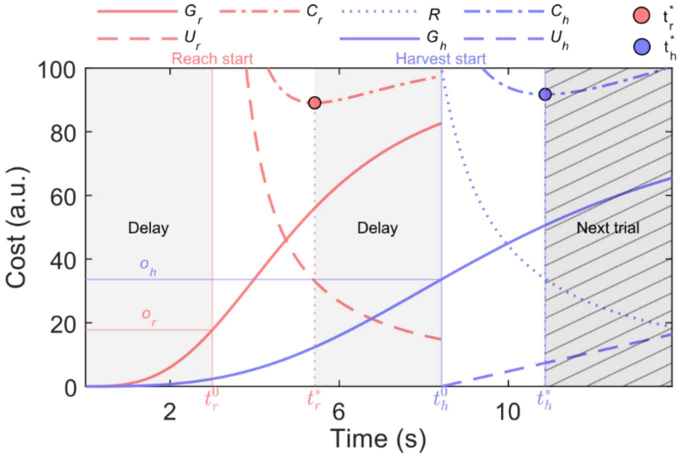


**Figure S6. Separate-cost model applied to the experimental paradigm.** The specific case is considered here where the sequence of phases is as follows: first delay, reach, second delay, harvest. Both reach and harvest time costs (red and blue solid lines, respectively) start at $\boldsymbol{t = 0}$ (supposedly the beginning of the trial). At the end of the first delay, the reach phase start with a cost of time already increased due to the time offset $o_r$, thereby affecting reach duration. Once the target has been reached ($\boldsymbol{t = t_r^*}$), the second delay starts, followed by the harvest phase. Because harvest starts after reaching (within a trial), reach phase duration is included in the harvest time offset in addition to both delays ($\boldsymbol{o_h = t_d + t_r}$), thereby affecting the harvest's time cost and influencing harvest duration. Consequently, the separate-cost model implies a possible inter-dependence between movement and decision, allowing co-regulation, depending on their sequential order. $U_r$: reach's cost of effort; $o_r$: reach's time offset; $G_r$: reach's cost of time; $C_r$: reach's total cost; $U_h$: harvest's cost of effort; $o_h$: harvest's time offset; $G_h$: harvest's cost of time; $R$: harvest's cost of reward $C_h$ : harvest's total cost; $t_h$ : harvest duration; $t_r$ : reach duration.

**Supplementary Video**

First, this video presents the predictions of the common-utility model. It begins with a general overview of how the global capture rate (J) depends on reach and harvest durations, with a maximum observed at the optimal durations ($t_h^*, t_r^*$). Next, simulations are presented showing how these optimal durations gradually change following an increase in reach effort (HRE, $t_{video} = 3\ s$), a decrease in reach effort (LRE, $t_{video} = 7\ s$), an increase in delays (HD, $t_{video} = 11\ s$), and a decrease in delays (LD, $t_{video} = 16\ s$). The results show that, regardless of how time and effort are modulated, both reach and harvest durations change, leading to a rigid co-regulation of both vigor.

Secondly, the predictions of the separate-cost model are presented. Similarly, gradual changes in optimal reach and harvest durations are shown following an increase in reach effort (HRE, $t_{video} = 23\ s$), a decrease in reach effort (LRE, $t_{video} = 27\ s$), an increase in delays (HD, $t_{video} = 31\ s$), and a decrease in delays (LD, $t_{video} = 35\ s$). In this case, some modifications result in theoretical co-regulation (e.g. HRE, LRE). Changes in reach duration are comprised within the harvest temporal offset ($o_h$), integrated by the cost of time. Meanwhile, some conditions reveal a decoupling (e.g., HD, LD) where changes in delay after reach are not integrated within the reach temporal offset but only in the harvest temporal offset, thereby leading to asymmetric changes in phase duration. Hence, the separate-cost model allows for flexible regulation of reach and harvest vigor, while the common-utility model implies a rigid co-regulation of both.

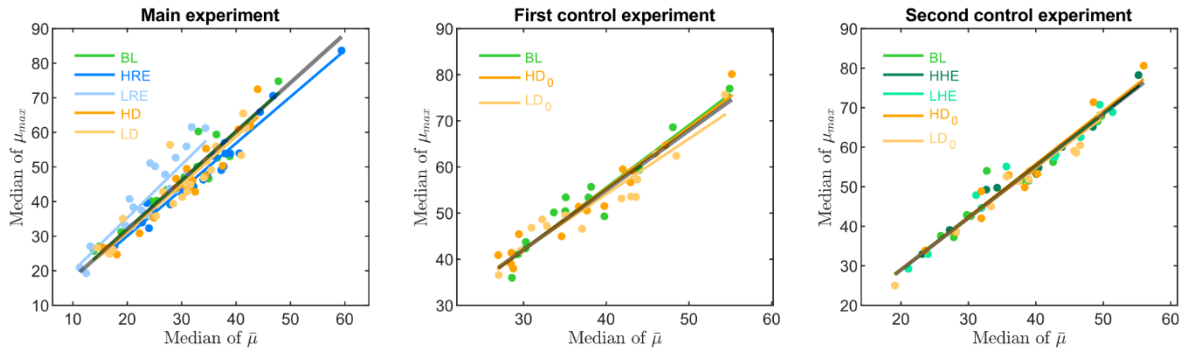


**Figure S7. Average normalized torque as a predictor of maximum normalized torque.** This figure depicts the relationship between the median of the average normalized torque and the median of the maximum normalized torque for each condition and individual, for the main experiment (left panel), the first control experiment (middle panel), and the second control experiment (right panel). Colored circles and lines represent respectively individual data points and regression line (random intercept and slope) for each condition. The black line represents the average regression with fixed intercept and slope. BL: Baseline; HRE: High Reach Effort; LRE: Low Reach Effort; HD: High Delay; LD: Low Delay; HHE: High Harvest Effort; LHE: Low Harvest Effort; $HD_0$: High Delay; $LD_0$: Low Delay.
